# IL-6 Trans-Signaling and Crosstalk Among Tumor, Muscle and Fat Mediate Pancreatic Cancer Cachexia

**DOI:** 10.1101/2020.09.16.300798

**Authors:** Joseph E. Rupert, Andrea Bonetto, Ashok Narasimhan, Yunlong Liu, Thomas M. O’Connell, Leonidas G. Koniaris, Teresa A. Zimmers

## Abstract

Most patients with pancreatic adenocarcinoma (PDAC) suffer unintentional weight loss, or cachexia. Interleukin-6 causes cachexia in mice and associates with mortality in PDAC. Here we show that tumor cell-derived IL-6 mediates crosstalk between tumor and peripheral tissues to promote cachexia. Tumor-cell IL-6 elicits expression of IL-6 in fat and IL-6 and IL-6 receptor (IL6R) in muscle, concomitantly raising both in blood. Inflammation-induced adipose lipolysis elevates circulating fatty acids, which cooperate with IL-6 to induce skeletal muscle dysmetabolism and wasting. Thus, PDAC induces crosstalk among tumor, fat and muscle via a feed-forward, IL-6 signaling loop. Tumor talks to muscle and fat through IL-6, and muscle to fat via IL6R trans-signaling, and fat to muscle through lipids and fatty acids. Disruption of this crosstalk by depletion of tumor-derived IL-6 halved fat wasting and abolished muscle loss, supporting IL-6, IL-6R and lipids as causal nodes for tissue crosstalk in PDAC cachexia.

**Significance:** PDAC-associated cachexia significantly increases patient morbidity and mortality. This study identifies muscle and fat crosstalk via IL6R trans-signaling in concert with muscle steatosis as a main driver of PDAC-associated cachexia.

## INTRODUCTION

Pancreatic ductal adenocarcinoma (PDAC) is one of the deadliest cancers with a five-year mortality rate of >91% (Siegel et al., 2019). Cachexia, the involuntary loss of fat, muscle and bone mass, affects over 80% of patients with PDAC and leads to increased morbidity and mortality (Hendifar et al., 2018; Sun et al., 2015; von Haehling et al., 2016). Both cancer and cachexia are associated with systemic inflammation affecting multiple organ systems (Argiles et al., 2018; Onesti and Guttridge, 2014). While various cytokines, chemokines, and growth factors are changed in PDAC, Interleukin-6 (IL-6) specifically has been positively correlated with PDAC presence (Holmer et al., 2014), disease progression (Ramsey et al., 2019), mortality (Babic et al., 2018; Suh et al., 2013), and cachexia (Okada et al., 1998) (Ebrahimi et al., 2004) (Martignoni et al., 2005). Although circulating IL-6 levels are not always detectable in early PDAC nor always correlated with cachexia severity (Ramsey et al., 2019; Talbert et al., 2018), higher tumor staining for IL-6 is associated with PDAC cachexia (Martignoni et al., 2005) and induction of monocyte IL-6 is predictive of survival in PDAC (Moses et al., 2009), suggesting that the serum levels of this short-lived cytokine might not be an appropriate measure of tissue activity. Functional data also support a role for IL-6 in PDAC tumor development (Lesina et al., 2011), progression (Zhang et al., 2013), metastasis (Razidlo et al., 2018), anti-tumor immunity (Flint et al., 2016), and response to chemotherapy (Long et al., 2017). IL-6 levels are high in PDAC models with weight loss (Flint et al., 2016) and IL-6 is functionally linked to cachexia in murine C26 colon adenocarcinoma and other models of cancer cachexia (Baltgalvis et al., 2008; Bonetto et al., 2012; Bonetto et al., 2011; Narsale and Carson, 2014). Moreover, IL-6 is sufficient to induce cachexia in mice (Baltgalvis et al., 2009; Chen et al., 2016; Tsujinaka et al., 1996) as well as lipolysis and atrophy in cultured adipocytes (Trujillo et al., 2004) and myotubes (Bonetto et al., 2012), respectively. IL-6 can be both detrimental and beneficial. While chronically increased IL-6 is associated with insulin resistance, inflammation, adipose tissue lipolysis, and muscle wasting in diseases from cancer and obesity to sepsis and burn injury (Kraakman et al., 2015; Pedroso et al., 2012; van Hall, 2012), acute expression of IL-6 promotes liver regeneration after injury (Jin et al., 2006; Koniaris et al., 2003) and is required for muscle regeneration, exercise-induced hypertrophy, and recovery from disuse atrophy (Begue et al., 2013; McKay et al., 2009) (Washington et al., 2011).

IL-6 initiates signal transduction by first binding to either the membrane-bound form of the interleukin-6 receptor (IL6R), also known as glycoprotein 80 (GP80), or its soluble form (sIL6R) (Schaper and Rose-John, 2015). Proteolytic shedding of a 55 kDa fragment in tissues expressing membrane IL6R results in circulating sIL6R (Schaper and Rose-John, 2015), an activity mediated in part through intracellular accumulation of phorbol esters and activation of protein kinase C theta (PKC-θ) (Mullberg et al., 1992). Both complexes of IL-6 with membrane or sIL6R bind the ubiquitously expressed membrane co-receptor IL-6 signal transducer (IL6ST), also known as glycoprotein 130 (GP130). The activity elicited by IL-6 and membrane IL6R is considered classical or cis signaling, while activity instigated by IL-6 with sIL6R is known as trans-signaling. Formation of either complex leads to trans-phosphorylation and activation of Janus kinases (JAKs), which phosphorylate the transcription factor Signal Transducer and Activator of Transcription 3 (STAT3), promoting STAT3 dimerization and translocation to the nucleus (Taniguchi and Karin, 2014). Unlike GP130, IL6R expression is not ubiquitous across cell types; thus, sIL6R trans-signaling allows for IL-6 signaling in IL6R negative cells (Rosean et al., 2014). Signaling through the membrane receptor is largely beneficial, while trans-signaling is generally pathological (Schaper and Rose-John, 2015). A growing number of studies suggest that neutralization of IL6R could have greater utility versus targeting IL-6 directly (Kraakman et al., 2015).

Given its various roles in PDAC, lipolysis and muscle wasting, here we investigated the effects of IL-6 emanating from PDAC tumor cells on tissue crosstalk and cachexia using isolated mouse PDAC cells deleted for IL-6 (KPC IL-6^KO^) in a mouse model of pancreatic cancer cachexia. Furthermore, we used this approach to characterize fat and muscle crosstalk via IL-6 signaling in pancreatic cancer cachexia. Our results suggest a feed-forward signaling loop between tumor-derived IL-6, skeletal muscle steatosis and sIL6R trans-signaling and adipose tissue IL-6 production working in concert to exacerbate skeletal muscle wasting in PDAC.

## RESULTS

### Human PDAC Tumor Cells are Heterogeneous for IL-6 Expression

Immunohistochemistry on human PDAC tumors obtained at surgery revealed stromal IL-6 staining in all tumors tested (Figure 1 A) as previously reported (Mace et al., 2018). However, IL-6 staining in tumor epithelial cells was also observed. Of 72 tumor samples, 32 demonstrated weak intensity IL-6 staining (Figure 1 A, top; and 1 B, left) and 40 samples strong intensity staining (Figure 1 A, bottom; and 1 B, right).

**Figure 1.**
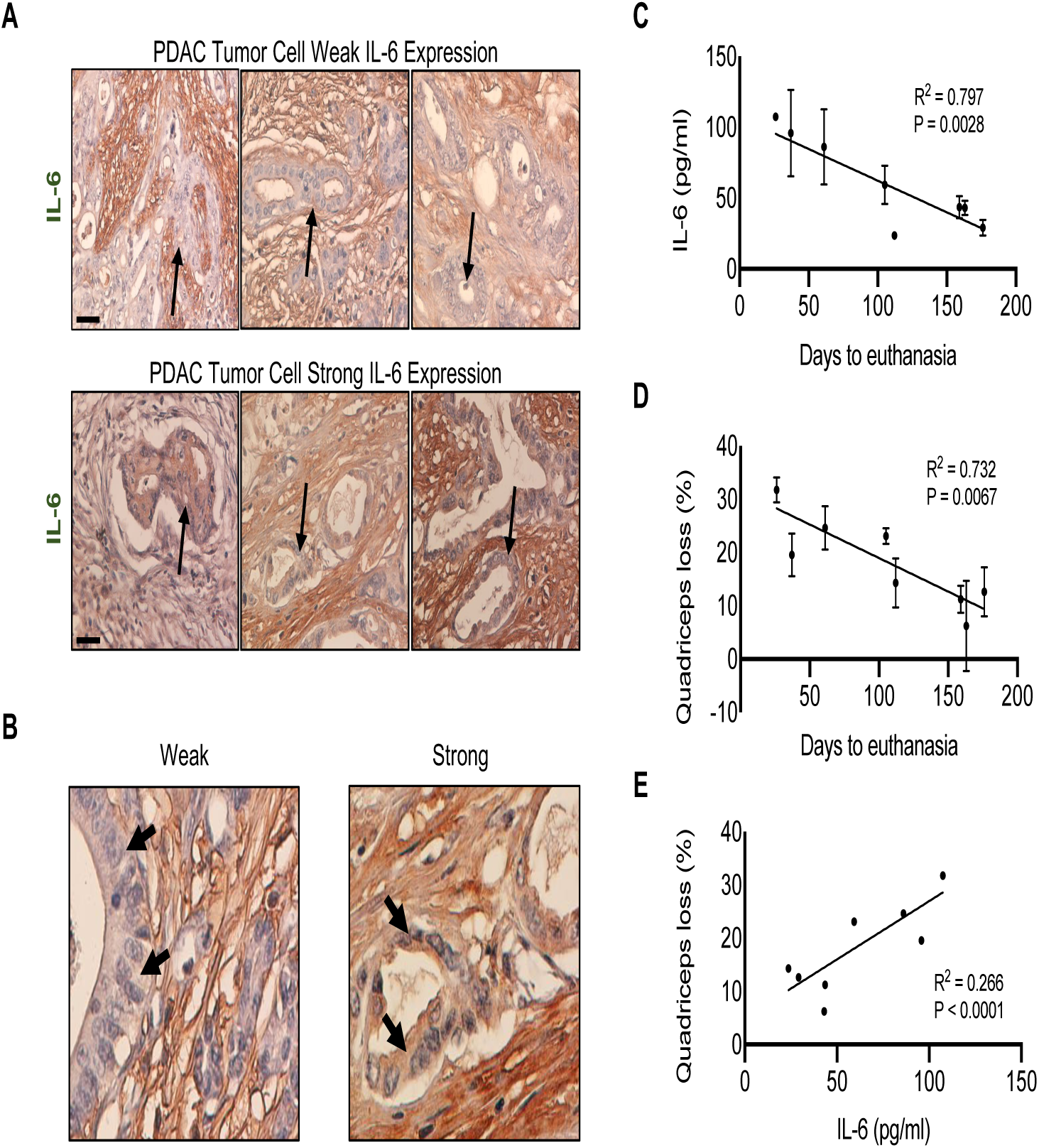
IL-6 protein expression is heterogeneous in human PDAC tumors and is associated with increased mortality and muscle wasting. Human PDAC tumors obtained were reacted for IL-6 immunohistochemistry (IHC). Tumor sections were classified as having either high or low expression of IL-6 specifically in tumor epithelial cells (arrows) (A). Of the 72 tumors, 40 had low tumor-cell IL-6 expression and 32 had high tumor-cell IL-6 expression. Images using increased magnification to show PDAC tumor cell IL-6 expression (B). Xenograft implantation of human tumors into mice showed correlations between human IL-6 and mortality (C), muscle loss and mortality (D), and human IL-6 and muscle loss (E). Scale bar = 40 μm.

Expression profiling of human and murine PDAC cell lines also confirmed a range of *IL6* expression from high (PSN-1, Panc03.27, PANC-1) to low (MIA PaCA, AsPC-1, HuPT4) (Figure S1). The importance of tumor cell-derived IL-6 was further investigated using human tumors in a murine xenograft model. Mice implanted with human PDAC tumors had a negative correlation when comparing human IL-6 expression in plasma and survival (Figure 1 C), loss of quadriceps mass and survival (Figure 1 D), and a positive correlation between loss of quadriceps mass and plasma IL-6 (Figure 1 E). These results indicate that PDAC epithelial cells are a source of IL-6 in the tumor micro- and macro environment and significantly contribute to cachexia and survival.

### PDAC-induced Cachexia and Mortality are Significantly Improved by Deletion of IL-6 from Tumor Cells

To evaluate the effects of tumor cell-derived IL-6 in PDAC cachexia we used CRISPR/Cas9 to edit the *Il6* gene in a cell line isolated from an autochthonic tumor arising in a C57BL/6 male KPC mouse. A similarly transfected clone not exposed to gRNAs was used as a control (KPC). *Il6* mRNA was detected in the parental KPC cell line (KPC-P) and the control KPC line, but not in the KPC-IL6^KO^ line (Figure 2 A); loss of expression was due to an insertional mutation (Figure S2). Both control and knockout lines exhibited similar growth characteristics in vitro (Figure 2 B), indicating that autocrine IL-6 was not required for proliferation.

**Figure 2.**
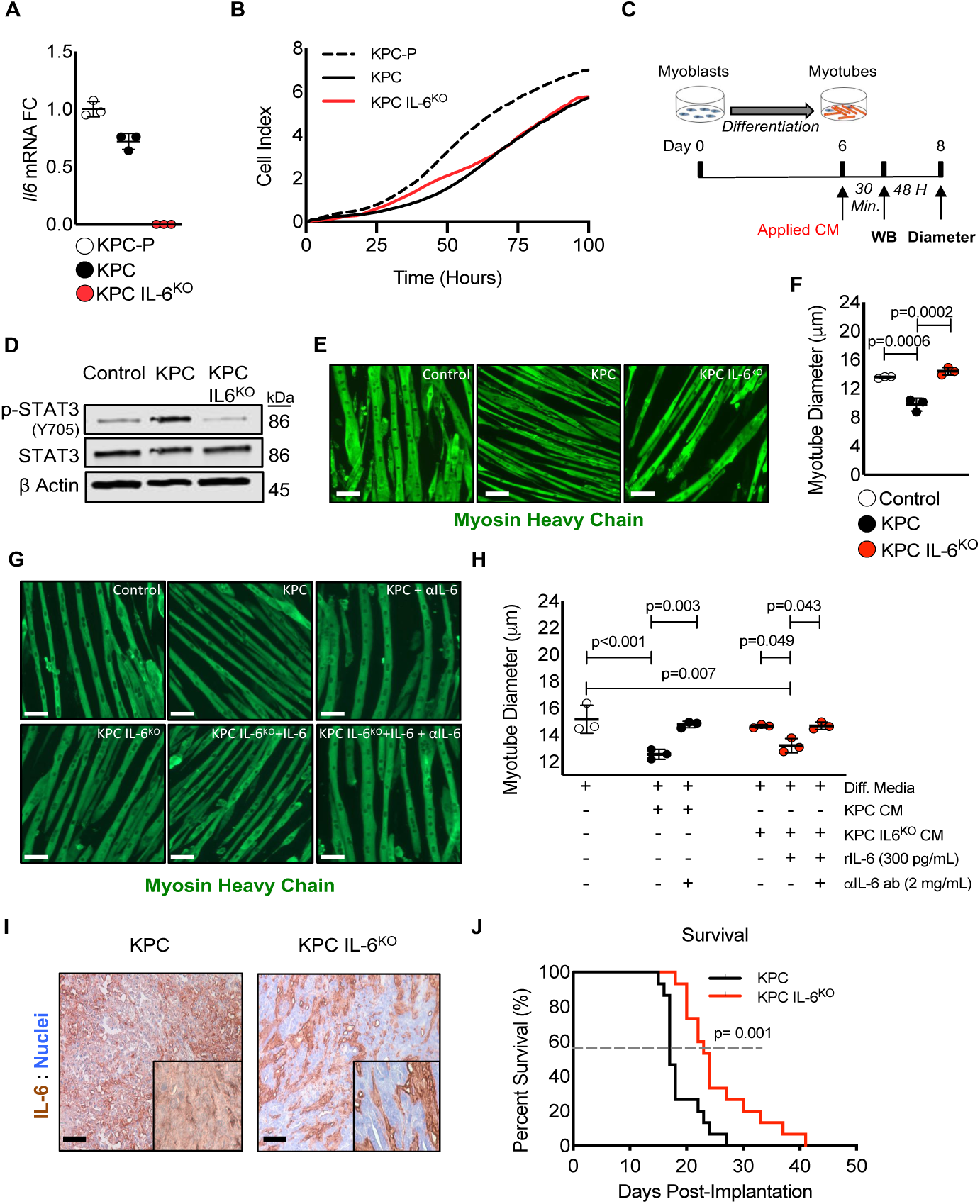
Deletion of IL-6 from KPC cells prevented muscle wasting in vitro and increased survival in mice. Targeted mutagenesis of the *Il6* gene was performed and a transfection control clone (KPC) and an *Il6* ablated clone (KPC IL-6^KO^) were selected for use in downstream experiments (**A**). The parental cell line (KPC-P) was used as a positive control for selection of clones (**A**). To determine if deletion of IL-6 affected tumor cell growth a proliferation assay was performed comparing the clones (**B**). Myoblasts were differentiated into myotubes and treated with 30% CM from clones to measure effects on myotube atrophy and STAT3 activation (**C**). Western blotting using myotube protein lysates were performed to measure STAT3 phosphorylation (**D**). Myotubes were visualized using florescent immunocytochemistry with an anti-myosin heavy chain antibody (**E**) and myotube atrophy measured (**F**). To verify atrophy was due to IL-6, myotubes were treated with KPC IL-6^KO^ CM plus recombinant IL-6 with and without the presence of an anti-IL-6 neutralizing antibody and the anti-IL-6 neutralizing antibody was also used to treat the KPC CM; myotubes were visualized with MHC IF and atrophy measured (**G and H**). KPC tumor cells were orthotopically implanted into mice, tumors were excised, sectioned and reacted with anti-IL-6 IHC to verify tumor cell IL-6 deletion (**I**). Survival was measured in mice orthotopically implanted with KPC and KPC IL-6^KO^ tumor cells (**J**). Scale bar = 20 μm; Error bars are standard deviation and significant differences are shown in the charts.

Deleting IL-6 also abolished muscle-wasting activity associated with KPC tumor cell conditioned media. Muscle growth and wasting is often modeled by manipulation or treatment of myotubes differentiated from C2C12 myoblasts (Figure 2 C). IL-6 itself induces atrophy of C2C12 myotubes via a STAT3-dependent mechanism (Bonetto et al., 2012). Here, conditioned media from KPC cells induced STAT3 phosphorylation (Figure 2 D). Myotubes were visualized using immunofluorescence for MHC (Figure 2E) and myotube diameter was measured resulting in a 25% reduction of myotube diameter in those treated with KPC media (Figure 2 F). An increase in STAT3 phosphorylation and a reduction in myotube diameter were absent in KPC-IL6^KO^CM treated myotube. To determine if the deletion of IL-6 from KPC IL-6^KO^ CM attenuating myotube atrophy, IL-6 was added back to KPC IL-6^KO^ CM media as well as neutralized in KPC CM and KPC IL-6^KO^ + IL-6 CM. Myotubes were again visualized using immunofluorescence for MHC (Figure 2 G) and diameters measured. The addition of IL-6 (300 pg/mL) to KPC IL-6^KO^ CM induced myotube atrophy (Figure 2 H), while the addition of IL-6 neutralizing antibody (2 mg/mL) to KPC CM and KPC IL-6^KO^ + IL-6 CM was sufficient to attenuate myotube atrophy (Figure 2 H). Next, the effects of deleting tumor cell-derived IL-6 in vivo were investigated. Orthotopic tumors from mice bearing KPC or KPC IL-6^KO^ tumors were reacted for IL-6 expression and positive IL-6 expression in KPC cells (Figure 2 I, left) and negative IL-6 expression in KPC IL-6^KO^ tumor cells (Figure 2 I, right) was confirmed. Finally, the deletion of tumor cell-derived IL-6 increased the survival of KPC IL-6^KO^ tumor mice compared to KPC tumor mice (Figure 2 J).

### Muscle Wasting is Spared in KPC IL-6^KO^ Tumor-bearing Mice

Apart from the survival study, two separate cohorts were analyzed for effects on cachexia, euthanizing mice when one group reached specified humane endpoints. Cachexia was prominent in the KPC group, with 11% to 26% wasting of limb muscles and heart (Figure 3 A), with significant loss of body weight (Figure S3) and loss of carcass mass (Figure S3). In contrast, muscle weights in the KPC-IL6^KO^ group were unchanged versus tumor-free control mice (Figure 3 A) with no change in body weight or carcass mass (Figure S3A and S3B). KPC tumor-bearing mice also had wasting of the liver (Figure S3C) and increased splenomegaly versus KPC IL-6^KO^ tumor-bearing mice (Figure S3D), although KPC IL-6^KO^ tumor-bearing mice had splenomegaly when compared to tumor-free mice (Figure S3D). This degree of wasting would be considered severe cachexia.

**Figure 3.**
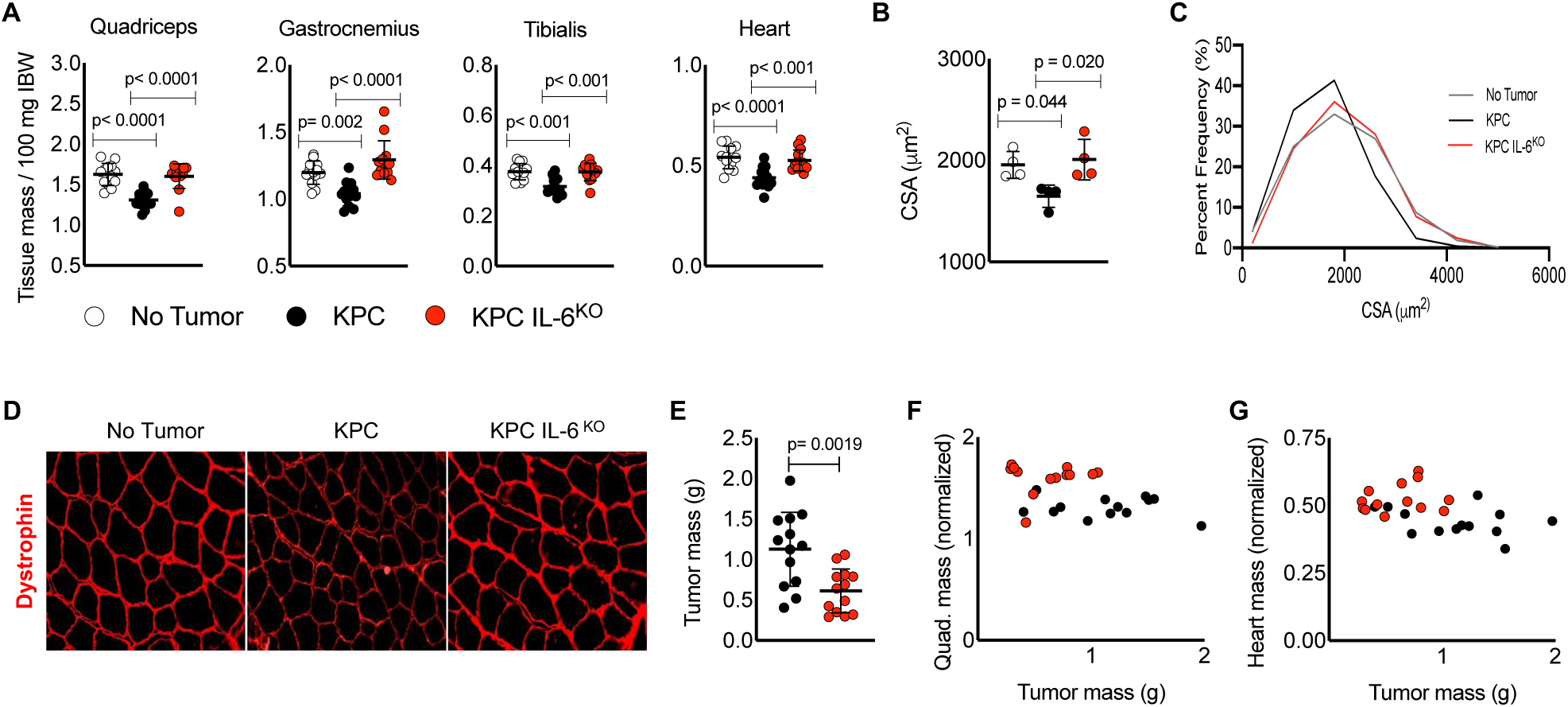
Deletion of tumor cell IL-6 attenuates muscle wasting. Mice were euthanized 17 days after injection due to severe cachexia in the KPC group and reaching humane endpoints. Skeletal muscles and the heart were excised at euthanasia and weighed and normalized to initial body weight (IBW) (**A**). Evaluation of muscle fiber CSA was done using sections from excised quadriceps muscles reacted for dystrophin expression (**D**) and mean fiber CSA measured (**B**). Cumulative fiber CSA values were organized based on percent distribution of fiber CSA and plotted to observe shifts in distribution (**C**). KPC IL-6^KO^ excised tumor mass was smaller than KPC tumor mass (**E**) thus, tumor mass was compared with muscle mass to determine if any correlation ns were present (**F and G**). Scale bar = 50 μm; Error bars are standard deviation and significant differences shown in charts.

The quadriceps muscles were sectioned and individual muscle fibers were visualized using immunofluorescence for dystrophin (Figure 3 B). Muscle from mice with KPC-IL6^KO^ tumors showed normal dystrophin expression, not the reduced expression described in murine cachexia (Acharyya et al., 2005; Stephens et al., 2015) and observed in mice with KPC tumors (Figure 3 B). Mice with KPC-IL6^KO^ tumors were also spared the myofiber atrophy observed in quadriceps of mice with control KPC tumors (Figure 3 C and 3 D). Mice bearing the KPC tumor had larger tumor mass than mice with KPC IL-6^KO^ tumors (Figure 3 E). Therefore, to investigate if tumor size was affecting muscle loss, a regression analysis was performed comparing tumor mass versus normalized muscle mass. There were no correlations between tumor mass and quadriceps mass (Figure 3 F) and tumor mass and heart mass (Figure 3 G). Tumor mass was also compared to the remaining skeletal muscles (Figure S3E and S3F) and epididymal fat pad mass (Figure S3G) with no correlations observed. To determine if deletion of IL-6 was playing a role in tumor grade and subsequently affecting the severity of cachexia, tumors were blindly scored in two independent evaluations. There was no difference in tumor grade between KPC and KPC IL-6^KO^ tumor-bearing mice (Figure S3H). Collectively, these results are consistent with IL-6 as a KPC-derived mediator of cachexia and mortality.

### Tumor Cell Deletion of IL6 Reduces Muscle Atrophy Pathway Activation

Muscle of mice with KPC-IL6^KO^ tumors also demonstrated reduced activation of atrophy pathways, including those involved in ubiquitin-proteasome mediated proteolysis, autophagy, translation repression, and mitochondrial function. Quadriceps muscles from mice with KPC tumors demonstrated increased total protein ubiquitination, as commonly observed in conditions of muscle catabolism, while total protein ubiquitination in mice with KPC IL-6^KO^ tumors was no different from controls (Figures 4 A and 4 B). Corroboratively, RNA levels for the atrophy-associated skeletal muscle E3 ubiquitin ligases *Fbxo32/Atrogin-1/MAFbx* and *Trim63/Murf1,* were only increased in KPC and not KPC-IL6^KO^ muscles (Figure 4 B, right). Autophagy and endoplasmic reticulum (ER) stress are also triggered in muscle atrophy marked by LC3B-II and Beclin-1, proteins with important roles in autophagosome formation (Sandri, 2016) and Atf4, which is increased by ER stress and the unfolded protein response (Bohnert et al., 2016). LC3B-II and ATF4 protein were increased in the muscle of KPC but not KPC IL-6^KO^ tumor-bearing mice (Figure 4 C and 4 D). While Beclin-1 was unchanged in muscle of mice with KPC tumors, it was actually decreased in muscle of mice with KPC IL-6^KO^ tumors (Figure 4 B and 4 D). Effects of these KPC tumors on protein synthesis markers in quadriceps were subtle, with no significant change in anabolic pAKT/AKT and p-mTOR/mTOR (Figure 4 E and 4 F). A significant decrease in p-4EBP1 in KPC muscle was suggestive of de-repression of translation (Figure 4 E and 4 F). There was no difference across groups in the master regulators of mitochondrial biogenesis, PGC1A and PGC1B (Figure 4 G and 4 H). While there was no difference in the mitochondrial uncoupling protein UCP2, an increase in UCP3 was observed in mice with KPC tumors, but not in those with KPC-IL6^KO^ tumors (Figure 4 G and 4 H). These results indicated that deletion of IL-6 from tumor cells largely abolished activation of muscle wasting pathways, especially those involved in the ubiquitin-proteasome and autophagy, in mice with PDAC tumors.

**Figure 4.**
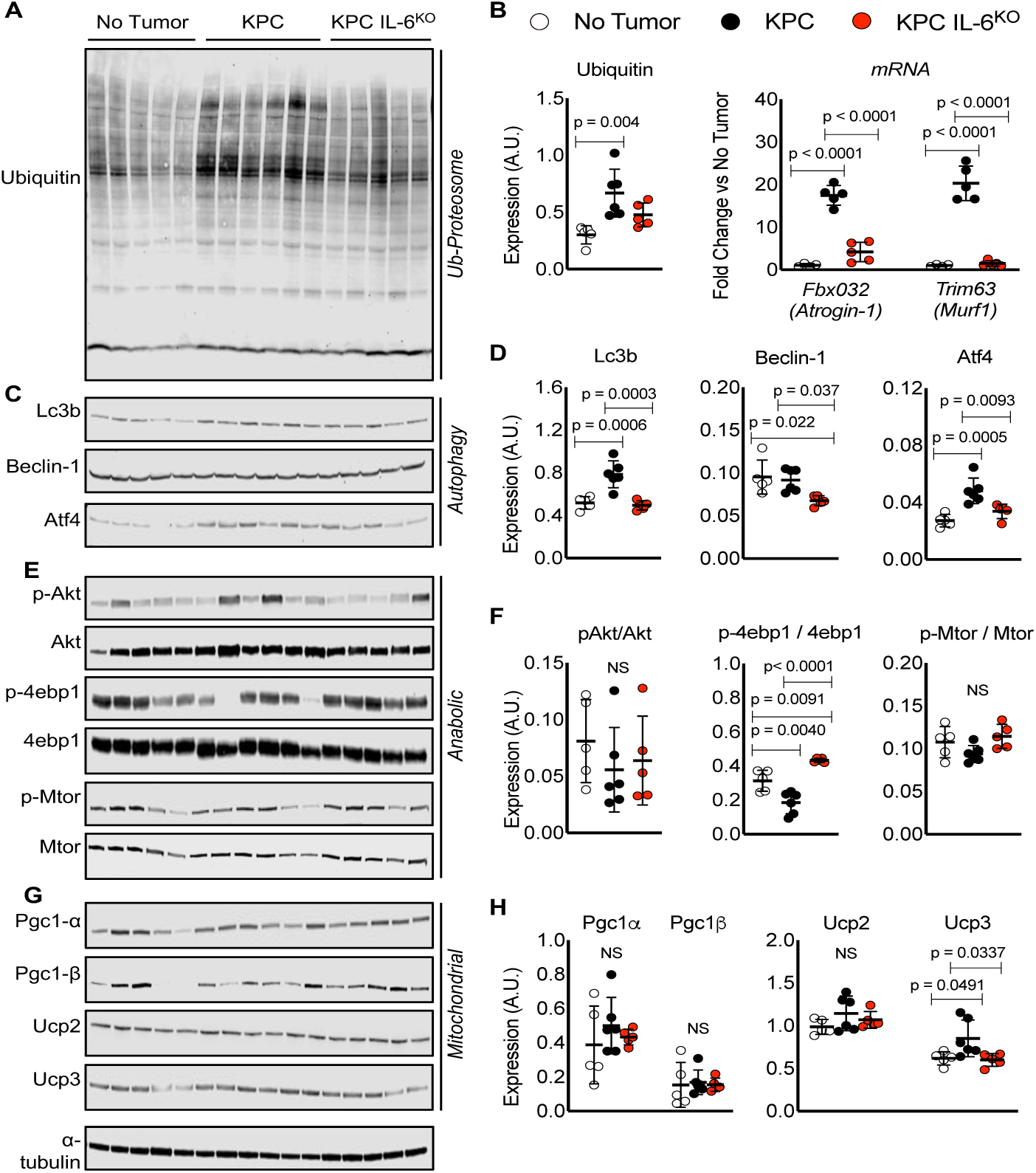
Measurement of protein expression for common molecular pathways associated with muscle wasting. Western blotting results evaluating muscle proteins involved in ubiquitination (**A and B**), autophagy (**C and D**), anabolism (**E and F**) and mitochondria biogenesis and metabolism (**G and H**) from protein lysates made from quadriceps harvested at euthanasia. Analysis of qPCR results for mRNA expression of E3 ubiquitin ligases *Atrogin-1* and *Murf1* in quadriceps was also performed (**B, right**). Error bars are standard deviation and significant differences between group means measured using ANOVA

### KPC but not KPC IL-6^KO^ Tumors Induced Inflammation, Lipid Accumulation and Oxidative Stress in Skeletal Muscle

RNA sequencing revealed 3480 up-regulated and 3793 down-regulated genes [fold change ≥ |1.5|; false discovery rate (FDR) of 0.05] in the muscle of KPC tumor mice, while only 24 up-regulated and 69 down-regulated genes were found in the muscle of mice with KPC-IL-6^KO^ tumors (Figure 5 A). Pathway analysis implicated a number of affected pathways in muscle including adipogenesis, oxidative stress, inflammation, fatty acid oxidation (all generally increased) and glycolysis, which was decreased (Figure 5 B-F). These were largely unchanged in mice with KPC-IL-6^KO^ tumors, save fatty acid oxidation, which was decreased relative to controls (Figure 5 E). Consistent with the pathway analysis, Oil Red O staining demonstrated intramyocellular lipid accumulation (myosteatosis) in quadriceps of KPC tumor-bearing mice, but not in KPC-IL-6^KO^ tumor-bearing or control mice (Figure 5 G, top; and 5 H, left). Succinate dehydrogenase (SDH) histochemistry, an indicator of mitochondrial respiration and localization, revealed aberrant localization and decreased respiratory capacity in the muscle of KPC tumor mice and to some degree in muscle from KPC IL-6^KO^ tumor mice (Figure 5 G, bottom; and 5 H, right).

**Figure 5.**
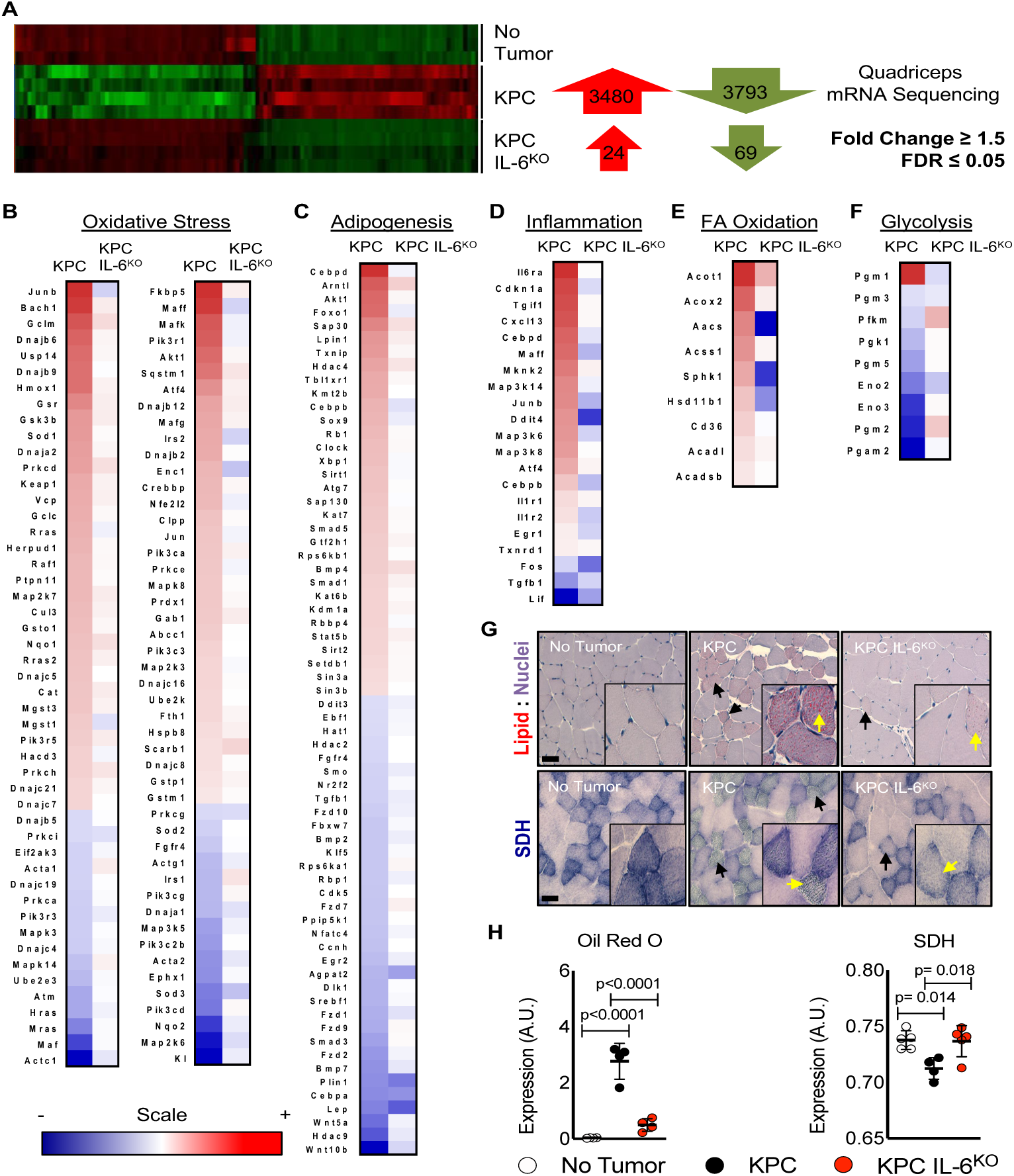
Deletion of IL-6 from KPC cells reduces activation of key cachexia pathways in muscle. Isolated RNA from the quadriceps of no tumor, KPC tumor and KPC IL-6KO tumor mice was sequenced and differentially regulated genes (fold-change ≥ 1.5 and FDR ≤ 0.05) were compared across groups (**A**). Ingenuity Pathway Analysis (IPA) using quadriceps RNA sequencing data identified various altered pathways and their associated genes (shown in heat map format) that have roles in muscle wasting including adipogenesis (**B**), Oxidative Stress (**C**), Inflammation (**D**), Fatty Acid (FA) Oxidation (**E**) and Glycolysis (**F**); the scale bar illustrates increased (Red) and decreased (Blue) gene expression for the heat maps. Measurements of muscle lipids using Oil Red O staining (**G and H**) and succinate dehydrogenase activity as a marker for mitochondria oxidative capacity (**G and I**) were performed on quadriceps muscle cross-sections; scale bar = 50 μm, arrows indicate fibers with increased lipid accumulation and aberrant SDH reactivity. Error bars are standard deviation and significant differences shown in charts

### Adipose Tissue is Not Preserved by Tumor-cell Deletion of IL-6

Because IL-6 has been shown to promote lipolysis, we evaluated tumor effects on adipose tissue wasting. Both KPC and KPC-IL-6^KO^ tumor mice had wasting of the epididymal fat pad versus no tumor mice, although fat wasting in KPC-IL-6^KO^ tumor mice was significantly attenuated (Figure 6 A). Consistent with this lesser fat wasting, plasma fatty acids and glycerol were increased only in the KPC tumor mice (Figure 6 B and 6 C). RNAseq of adipose tissue demonstrated a similar intermediate phenotype, with 268 up-regulated and 145 down-regulated genes in fat pads of mice with KPC tumors, and 209 up-regulated and 0 down-regulated genes in the KPC-IL-6^KO^ group (fold change ≥ |1.5|; FDR 0.05) (Figure 6 D). Granulocyte adhesion/diapedesis, LXR/RXR activation, acute phase signaling, and IL-6 signaling were among the top altered pathways associated with cachexia in adipose tissue (Figure 6 E-H), with activated genes similar across both tumor conditions. These results suggest that adipose tissue is more sensitive to tumor-produced factors as well as lower plasma IL-6 levels than muscle.

**Figure 6.**
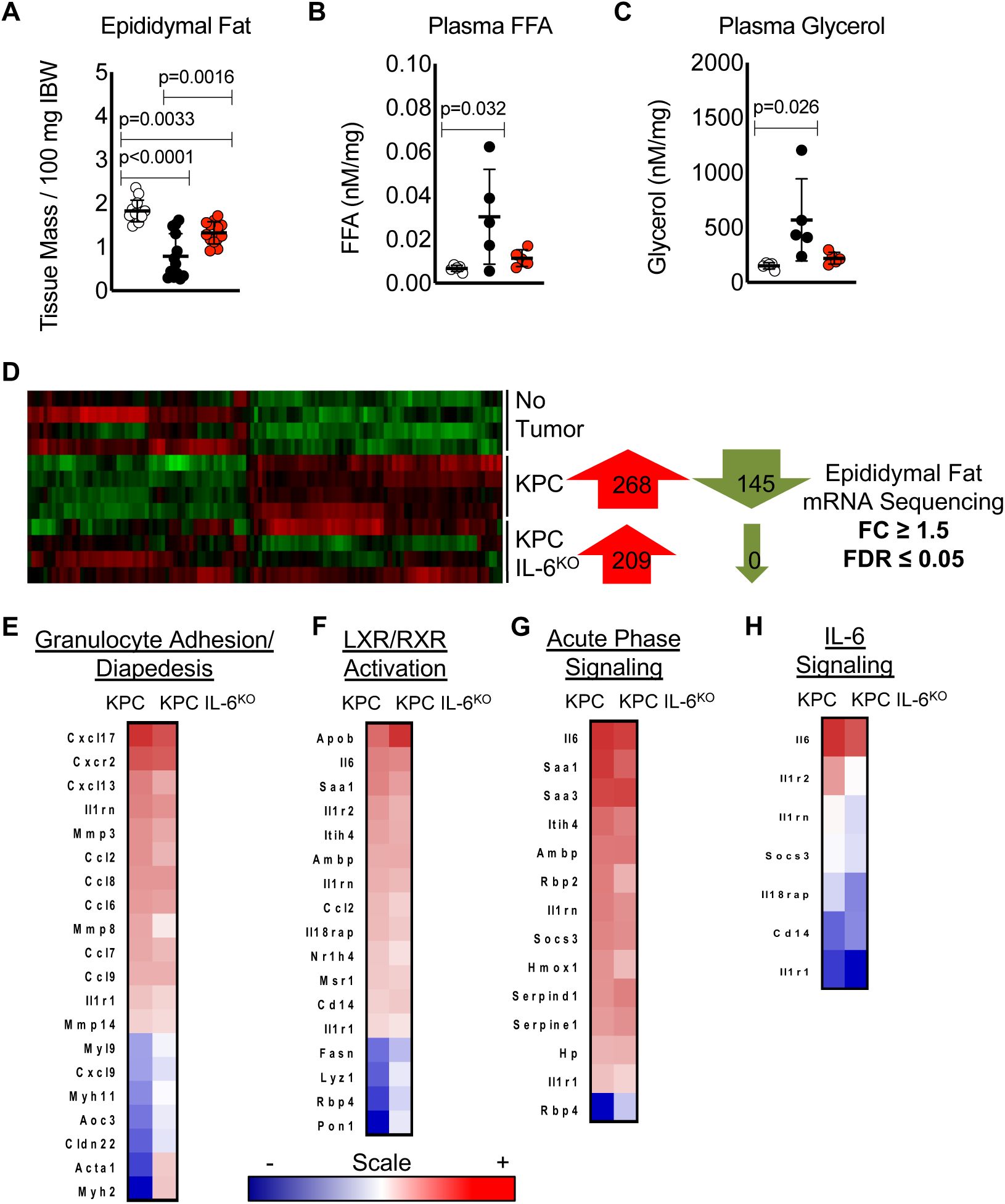
Deletion of IL-6 from KPC cells reduces fat wasting but did not hinder change in gene expression versus KPC tumor mice. Epididymal fat pad mass was measured at euthanasia and normalized to initial body weight (IBW) (**A**).Characterization of lipolysis using measurements of plasma glycerol (**B**) and fatty acids (**C**) normalized to epididymal fat pad mass from mice. Isolated RNA from the epididymal fat pads was sequenced and differentially regulated genes (compared to no tumor group, fold-change ≥ 1.5 and FDR ≤ 0.05) were compared across groups (**D**). Ingenuity Pathway Analysis (IPA) using epididymal fat pad RNA sequencing data identified various altered pathways and their associated genes (shown in heat map format) that have roles in inflammation and lipolysis including Granulocyte Adhesion/Diapedesis (**E**), LXR/RXR Activation (**F**), Acute Phase Signaling (**G**) and IL-6 Signaling (**H**); the scale bar illustrates increased (Red) and decreased (Blue) gene expression for the heat maps.

### IL-6 Pathway Proteins are Differentially Expressed in Fat Versus Muscle of Mice with PDAC, Implicating IL6R Trans-signaling from Muscle to Fat

Given the differential sensitivity of muscle and fat to tumor-derived IL-6, we sought to investigate tissue crosstalk in PDAC cachexia. As expected, implanted KPC tumor cells and host stromal cells expressed IL-6, while KPC-IL6^KO^ tumors showed expression only in stroma (Figure 2 I). Plasma levels of IL-6 were correspondingly increased, with a significant increase (∼150-fold) in KPC tumor mice over baseline levels, while mice with KPC-IL6^KO^ tumors showed roughly half that increase, and the difference were not significantly different from normal controls (Figure 7 A). Plasma soluble IL-6 receptor (sIL6R), which is largely produced by shedding of a 55 kDa fragment from the membrane-bound IL6R into the circulation, was also significantly increased in KPC tumor mice (∼25-fold) over controls; plasma sIL6R levels in KPC-IL6^KO^ mice were not different from controls (Figure 7 B).

**Figure 7.**
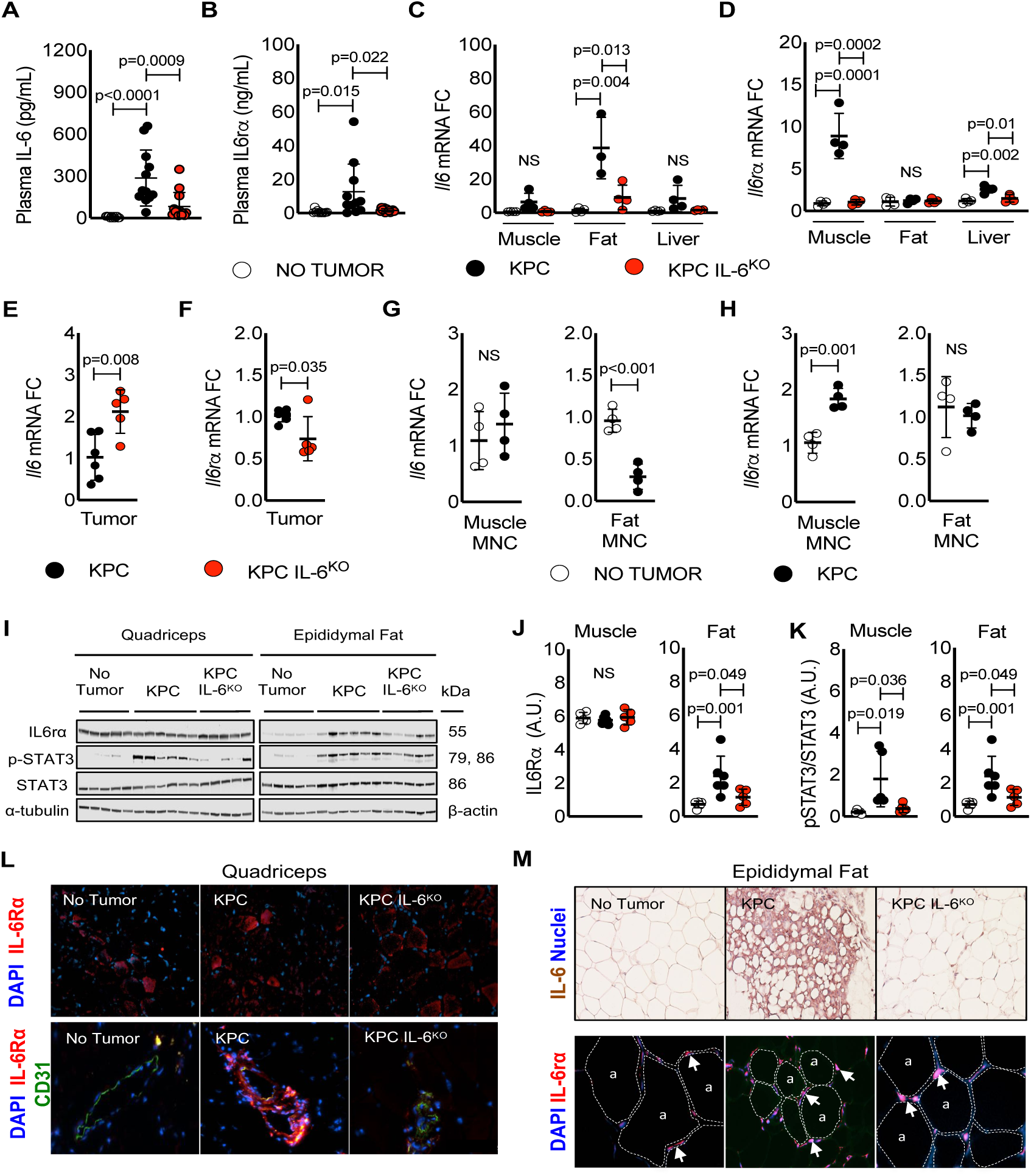
Evidence for an IL-6, IL6R circuit among tumor, adipose tissue and skeletal muscle in PDAC cachexia. Plasma from no tumor, KPC tumor and KPC IL-6^KO^ tumor mice was harvested immediately prior to euthanasia and measured for IL-6 (**A**) and IL6R (**B**) protein expression using an ELISA. Isolated RNA from quadriceps, epididymal fat, and liver of mice was used to measure mRNA expression of *Il6* (**C**) and *Il6r* (**D**) in tissues and presented as fold change versus no tumor mice. Isolated RNA from KPC and KPC IL-6^KO^ tumors was used to measure mRNA expression of *Il6* (**E**) and *Il6r* (**F**) in tumors and presented as fold change versus KPC tumors. To determine with increased detail the source of *Il6* mRNA in fat and *Il6r* in muscle, tissues were dissociated and isolated RNA from the mononuclear cell fractions (MNC) was used to measure mRNA expression of *Il6* (**G**) and *Il6r* (**H**). Protein expression for IL6R and STAT3 phosphorylation was quantified in the quadriceps and epididymal fat pads of mice using western blotting (**I, J, K**). Immunofluorescence showing the expression of IL6R protein (red) and nuclei (blue, DAPI) in the quadriceps muscle fibers of mice (**L, top**) and IL6R protein (red), the endothelial protein marker CD31 (green) and nuclei (blue, DAPI) to visualize IL6R expression in and around blood vessels in the quadriceps muscle of mice (**L, bottom**). IHC for IL-6 protein expression in the epididymal fat pads of mice (**M, top**) and immunofluorescence showing the expression of IL6R protein (red) and nuclei (blue, DAPI) in epidiymal fat (**M, bottom**); arrows indicate positive IL6R staining and dotted lines outline adipocytes denoted with the letter “a”. Error bars are standard deviation and significant differences between group means are shown in the charts.

To identify the sources of IL-6 and sIL6R, we measured *Il6* and *Il6r* mRNA expression in muscle, fat, liver and tumor and protein expression for IL6R and phosphorylation of STAT3 in the quadriceps muscle and the epididymal fat pads via western blotting. The results indicate that both muscle and fat express IL-6 in PDAC cachexia, and further that muscle is a source of circulating sIL6R, enabling trans-signaling of IL-6 in adipose tissue. Specifically, *Il6* mRNA was significantly increased 37-fold in epididymal fat from mice with KPC tumors (Figure 7 C) and although not significant likely from increased variation and smaller sample size, a trend in increased *Il6* was observed in muscle (6.4-fold, p=0.06) and liver (8.5-fold, p=0.1) from KPC tumor-bearing mice (Figure 7 C). In contrast, *Il6r* mRNA was significantly increased in the quadriceps (8.9-fold) and liver (2.6-fold) but unchanged in the adipose tissue from KPC tumor mice (Figure 7 D). Interestingly, compared to KPC tumors, *Il6* mRNA was increased in the KPC IL-6^KO^ tumors (Figure 7 E), yet tumor *Il6r* mRNA was actually decreased in KPC IL-6^KO^ tumors (Figure 7 F). Since changes in *Il6* and *Il6r* mRNA expression were only observed in muscle and fat of KPC tumor-bearing mice, we investigated in more detail the source of these altered gene expressions by dissociating the tissue and analyzing the mononuclear cell fractions (MNC) in no tumor and KPC tumor-bearing mice. Compared to no tumor mice, *Il6* mRNA expression was unchanged in the muscle MNC fraction from KPC tumor-bearing mice while unexpectedly; it was reduced in the fat MNC fraction from KPC tumor-bearing mice (Figure 7 G). *Il6r* mRNA expression was increased, although modestly, in the muscle MNC fraction from KPC tumor-bearing mice and unchanged in fat compared to no tumor mice (Figure 7 H). These results indicate that the MNC fractions from muscle and fat are not significantly contributing to *Il6* mRNA expression, but the muscle MNC fraction likely contributes in part to whole tissue *Il6r* mRNA expression in skeletal muscle.

In contrast to the mRNA expression, there was no change in IL-6R protein expression in quadriceps and a significant increase of IL-6R protein in adipose tissue of KPC tumor mice (Figure 7 I and 7 J). Interestingly, only KPC mice had significantly increased STAT3 phosphorylation (pSTAT3) in quadriceps, while both tumor-bearing groups had increased pSTAT3 in fat (Figure 7 I and 7 K). Immunofluorescence demonstrated IL6R protein in myofibers in all groups (Figure 7 L, top) as well as robust staining in muscle blood vessels only in the KPC tumor mice (Figure 7 L, bottom). Strong IL-6 expression was observed in the fat from KPC tumor-bearing mice (Figure 7 M, top). In adipose tissue, visualization of the IL6R showed accumulation near cells positioned between adipocytes in all groups (Figure 7 M, bottom). Taken together, these results suggest tumor cell and adipose tissue IL-6 production contribute to systemic IL-6 levels and intricate crosstalk between stromal and tumor cells may be occurring in the KPC IL-6^KO^ tumors requiring further investigation. Furthermore, these findings also implicate muscle as a primary source for circulating sIL6R, which likely promotes its accumulation in fat.

### Similar Changes to Tissue Wasting and Il6 and Il6r mRNA Expression are Observed with In Vitro Studies of the Tumor-Adipose-Muscle Crosstalk

Our results indicate that KPC tumors increase lipid accumulation, *Il6r* expression, and atrophy in skeletal muscle, and also activate lipolysis and *Il6* expression in fat (Figures 5-7). These effects on distant tissues could be mediated directly by tumor-derived products or indirectly through other cellular or molecular mediators. I tested whether products of KPC cells could affect muscle and fat directly. Indeed, KPC conditioned media (CM) was associated with myotube atrophy (Figure 8 A) and similar to in vivo results, increased *Il6* and *Il6ra* mRNA expression was observed in myotubes after KPC CM treatment (Figure 8 B). KPC CM also increased 3T3L1 adipocyte lipolysis as measured by media glycerol concentration (Figure 8 C) and again, in parallel with our in vivo results, *Il6* but not *Il6ra* mRNA expression was increased in 3T3L1 adipocytes after treatment with KPC CM (Figure 8 D). KPC tumor-induced fat wasting produced increased circulating fatty acids and glycerol in vivo (Figure 6 B and 6 C).

**Figure 8.**
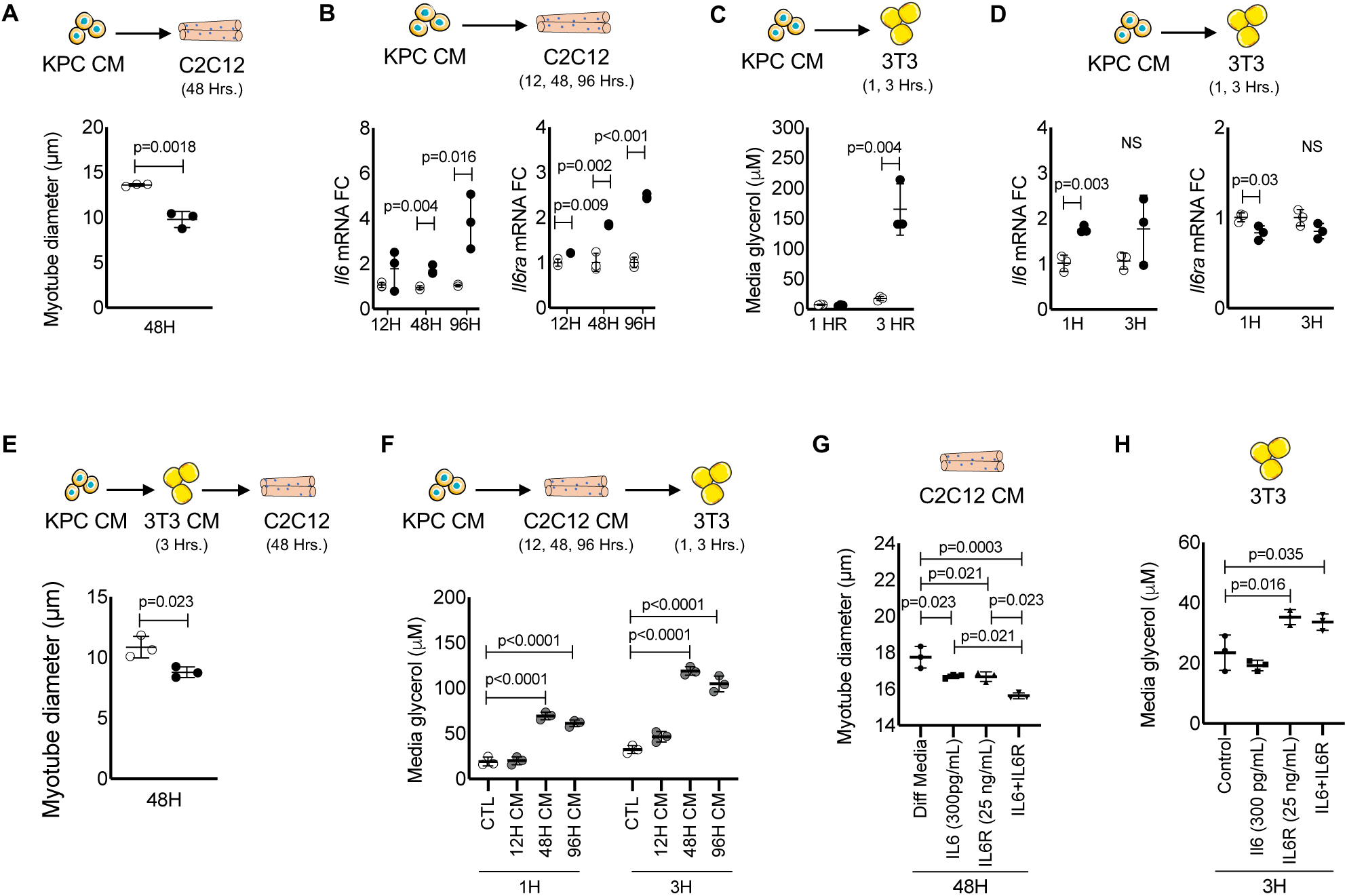
Modeling of IL-6, IL6R tumor-tissue crosstalk in vitro. C2C12 myotubes were incubated with conditioned media (CM) from KPC tumor cells for 48 hours and atrophy measured (**A**) and for 12, 24 and 96 hours and isolated RNA was used to measure mRNA expression of *Il6* and Il6r at each time point (**B**). 3T3 adipocytes were incubated with KPC CM for one and three hours and glycerol was measured as a marker for lipolysis (**C**); 3T3 adipocyte RNA was harvested at each time point and used to measure mRNA expression of *Il6* and *Il6r* (**D**). The tumor-fat-muscle crosstalk was investigated by incubating 3T3 adipocytes with KPC CM and then using the 3T3 CM to treat myotubes for 48 hours and measure atrophy (**E**). The tumor-muscle-fat crosstalk was investigated by treating myotubes for 12, 24 and 96 hours with KPC CM and then using the myotube CM to treat 3T3 adipocytes for 1 and 3 hours to measure lipolysis via media glycerol content (**F**). To decipher the effects of IL-6 and IL-6R in muscle and fat, myotubes and adipocytes were treated in vitro. Myotubes were incubated with differentiation media (DM), physiological levels of recombinant murine IL-6, physiological levels of recombinant murine IL6R, or the combination of IL-6 and IL6R for 48 hours and atrophy measured (**G**). 3T3 adipocytes were incubated with growth media (Control), physiological levels of recombinant murine IL-6, physiological levels of recombinant murine IL6R, or the combination of IL-6 and IL6R for 3 hours and media glycerol measured (**H**). Significant differences are shown in each chart.

Moreover, increased *Il6r* mRNA expression was increased in muscle but IL6R protein was increased in fat. Extracellular lipids have been implicated in muscle dysfunction in diabetes and metabolic syndrome, and muscle IL6Rexpression, but the effects in PDAC cachexia are not described. We modeled the effect of the tumor-fat-muscle crosstalk axis in vitro using a series of CM swapping experiments. 3T3L1 adipocytes were treated using KPC CM to induce lipolysis and the resulting 3T3L1 CM was then used to treat myotubes and measure the effects on myotube diameter. Myotubes treated with CM from 3T3L1 adipocytes after KPC CM treatment showed significant decrease in myotube diameter (Figure 8 E). To investigate the tumor-muscle-fat crosstalk axis, we treated C2C12 myotubes with KPC CM to induce atrophy and used the resulting C2C12 CM to treat 3T3L1 adipocytes. Treatment of 3T3L1 adipocytes with C2C12 myotube CM after KPC CM treatment significantly induced lipolysis, which was determined by glycerol release into the media (Figure 8 F). Finally, we investigated whether physiological concentrations of IL-6, IL-6R or the combination could induce wasting in both myotubes and adipocytes. Myotube diameter was decreased in the presence of IL-6 (300 pg/mL) or IL6R (25 ng/mL) versus controls, however, the combination of IL-6 and IL6R produced the most severe myotube atrophy (Figure 8 G). Interestingly, IL-6 treatment of 3T3L1 adipocytes using physiological levels did not induce lipolysis, yet presence of the IL6R was sufficient to induce lipolysis with or without exogenous IL-6 (Figure 8 H). These results suggest that adipocytes are likely more sensitive to lipolysis in the presence of the IL6R and not IL-6 alone. These findings also support that IL-6 and lipids can both mediate muscle loss, and further, that activation of adipose lipolysis likely contributes to muscle wasting in PDAC.

## DISCUSSION

Here we demonstrate novel tumor-tissue crosstalk in the macroenvironment of PDAC cachexia and further provide evidence of central roles for tumor cell-derived IL-6 and IL-6 trans-signaling. IL-6 was produced by both tumor cells and stromal cells in the tumor microenvironment; this signal was amplified distantly via by production of IL-6 from adipose tissue and skeletal muscle in the “tumor macroenvironment” of the whole host body. Adipose tissue loss was proportionate to circulating IL-6 and halved by elimination of IL*-*6 from tumor cells, while muscle loss was abolished by tumor-cell depletion of IL-6. These results indicate that adipose tissue was proportionately more sensitive to the effects of PDAC as measured by the induction of adipose wasting, lipolysis, and release of fatty acids and glycerol into the blood. Under these conditions, skeletal muscle exhibited steatosis, mitochondrial dysfunction, metabolic impairment, and wasting. As well, *Il6r* mRNA expression was increased by PDAC only in muscle and not fat, yet soluble IL6R protein accumulated in fat, and STAT3 phosphorylation indicated signaling in both peripheral tissue compartments. These results suggest that sIL6R is produced outside adipose tissue but accumulates in fat to mediate IL-6 signaling on adipocytes and further, that skeletal muscle is a major source of this sIL6R in cachexia. We also demonstrate that products of lipolysis (modeled with palmitate) exacerbate muscle wasting induced by IL-6, providing an explanation of how adipose loss can lead to muscle wasting. Thus, PDAC induces a feed-forward, IL-6 signaling loop and cross-talk among tumor, adipose tissue, and skeletal muscle, whereby tumor talks to muscle and adipose through IL-6, and muscle to adipose through IL6R, and adipose to muscle through palmitate and other fatty acids (Figure 9). Disruption of this signaling loop via depletion of IL-6 from tumor cells was sufficient to halve adipose wasting and abolish muscle wasting, supporting IL-6 production and signaling as causal nodes in PDAC cachexia.

**Figure 9.**
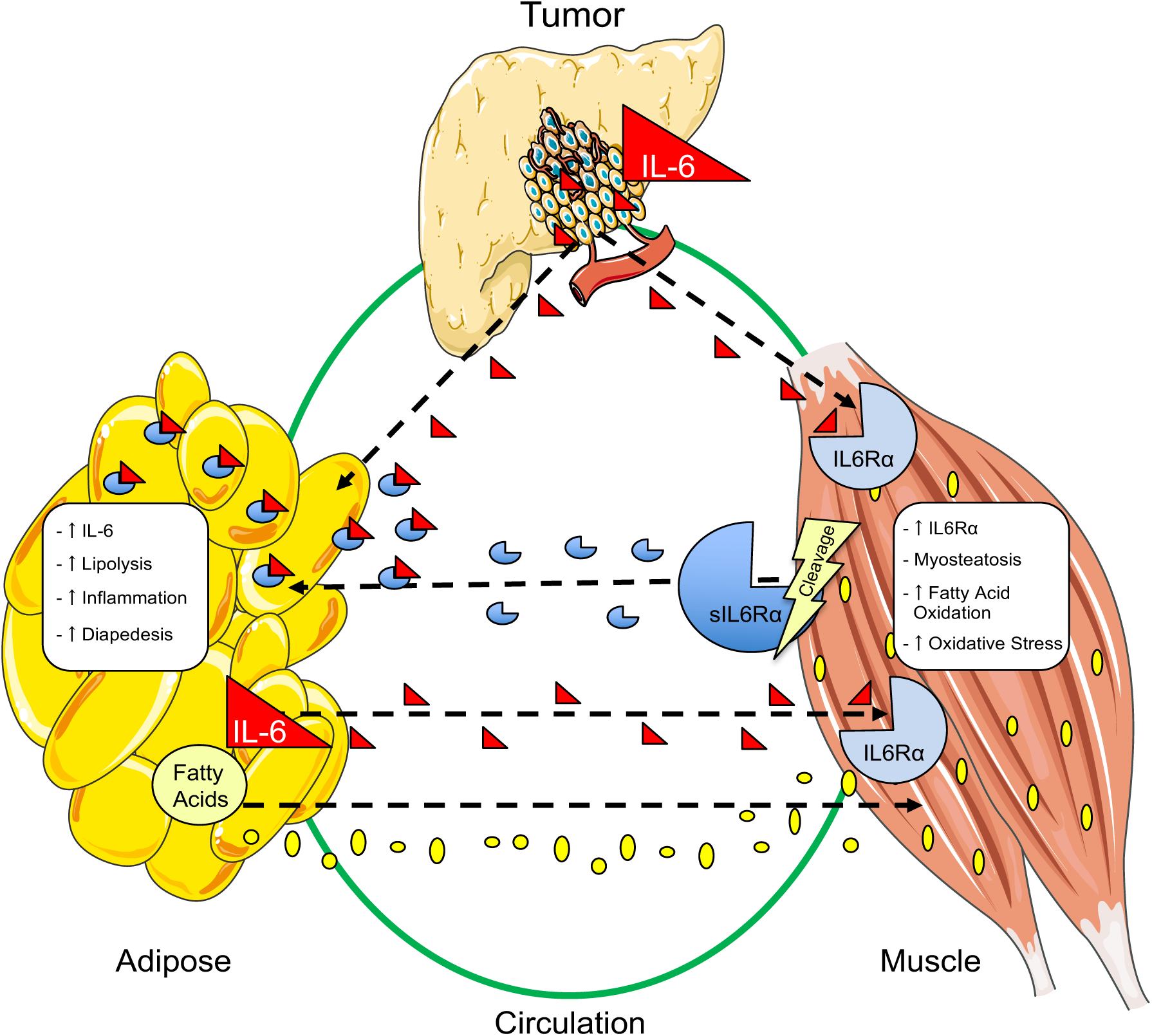
Illustration of tumor-fat-muscle crosstalk in PDAC. Using my results this illustration summarizes my findings with respec to tumor, fat and muscle crosstalk in PDAC.

Blood levels of IL-6 are generally correlated with increased cachexia and mortality in patients with late stage PDAC, and IL-6 production by fibroblasts and myeloid cells in the tumor microenvironment has been implicated in PDAC development and tumor growth, progression, metastasis and drug resistance (Holmer et al., 2014; Mace et al., 2018; Ohlund et al., 2017). However, IL-6 production by tumor epithelial cells is largely unstudied in PDAC. Our data indicate that a substantial number of human PDAC tumors exhibit expression of IL-6 in the epithelial compartment and further, that IL-6 is produced by a majority of established PDAC tumor cell lines. This has clear implications for PDAC tumor biology. In our studies, autocrine IL-6 production was not required for KPC tumor cell proliferation in vitro, although IL-6 null tumors in vivo trended smaller. Mice with IL-6 null tumors also exhibited reduced circulating IL-6, lesser adipose wasting and no muscle wasting. These phenomena are likely linked, although precisely how is unclear. Whether changes in tumor growth were due to IL-6-mediated differences in tumor phenotype or to reduced substrate availability from the relative preservation of fat and muscle, and conversely, whether reduced cachexia was due to decreased circulating IL-6 or also to other IL-6-dependent tumor properties including altered tumor phenotype and secretome are complex questions under active investigation in our laboratory. Regardless, clearly IL-6 production from tumor cells *can* orchestrate cachexia and this tumor characteristic could have diagnostic and therapeutic implications for PDAC.

How does tumor-derived IL-6 modulate cachexia? This study suggests that PDAC tumors elicit adipose tissue inflammation and lipolysis that in in turn induces muscle wasting, and that IL-6 is an important component of both. In the presence of tumor-cell IL-6, both muscle and fat mass were lost, and tissue gene expression profiles were massively changed from normal. However, in the absence of tumor-cell IL-6, only adipose tissue was greatly changed, both in mass and gene expression. Skeletal muscle was mostly normal in mice with IL-6 null tumors. This indicates that adipose tissue is more sensitive than muscle to the presence of tumor. Adipose inflammation is noted in patients with cancer cachexia (Camargo et al., 2015), and furthermore, as in our mice, the magnitude of fat loss (∼30%) exceeded that of muscle loss (7%) in patients on FOLFIRINOX for advanced disease (Kays et al., 2018), while only fat was lost in patients with PDAC on neoadjuvant therapy (Sandini et al., 2018).

Low skeletal muscle mass is predictive of mortality across diseases (Martin et al., 2013), which could be attributable to functional decline and respiratory or cardiac failure; however, the significance of fat loss is less clear. Indeed, the authors of a recent study discount adipose loss as participating in PDAC mortality, based upon feeding pancreatic enzymes to mice with pancreatic cancer and upon cross sectional body composition data in patients (Danai et al., 2018). Against that singular interpretation, however, are both clinical-correlative and mechanistic studies. Fat loss in the absence of muscle loss has been observed in cohorts of patients with PDAC (Sandini et al., 2018), (Kays et al., 2018), consistent with these being separable phenomena, and higher rates of adipose loss are associated with mortality in PDAC (Di Sebastiano et al., 2013). As well, obese patients with PDAC exhibit higher losses in weight, skeletal muscle, and adipose tissue along with poorer survival (Dalal et al., 2012). Even absent muscle wasting, low fat mass or “adipopenia” is associated with mortality in patients with diffuse large B-cell lymphoma treated with immunotherapy (Camus et al., 2014) and in patients with heart failure (Melenovsky et al., 2013), among other conditions. Furthermore, our longitudinal analysis of patients with advanced PDAC on FOLFIRINOX demonstrates equivalent mortality for patients with fat-only loss versus patients with both fat and muscle loss, with overall 10 months reduced survival than patients without loss of either depot (Kays et al., 2018). Thus, the preponderance of clinical data point to an important role for adipose in PDAC cachexia and mortality. Moreover, preserving adipose tissue via pharmacological or genetic manipulation of lipolysis and lipogenesis protects muscle in other cachexia models (Das et al., 2011; Rohm et al., 2016) (Tsoli et al., 2016), although in all of these studies multiple tissues in addition to adipose were affected by the intervention.

We posit that adipose wasting causes muscle wasting and thus contributes to mortality. Lipolysis of adipose tissue results in high circulating fatty acids and subsequent lipid uptake and accumulation in muscle, which after a period of metabolic adaptation ultimately results in lipotoxicity, including metabolic derangement, cellular stress, and muscle wasting. Our histological and RNAseq data support this interpretation. RNA sequencing analysis revealed genes associated with IL-6 signaling, lipid metabolism, and glycolysis in muscle. In the epididymal fat pads, pathways related to diapedesis, LXR/RXR activation—important regulators of cholesterol, lipid and glucose homeostatis, acute phase signaling, and IL-6 signaling pathways were affected. Importantly, gene expression correlated with wasting in both tissues. While muscle from mice with IL-6 null tumors was unaffected for mass and nearly normal in terms of gene expression, adipose wasting and gene expression changes were clearly well established even in the absence of tumor-cell IL-6. Tissue wasting also correlated with STAT3 phosphorylation, which was increased in the fat of both KPC tumor bearing mice as well as KPC-IL6ko tumor-bearing mice but only increased in muscle of KPC tumor mice. Furthermore, we only observed increased lipid accumulation in the muscle of KPC tumor mice and not in muscle from IL-6 null tumor mice. Impaired beta-oxidation is central to muscle wasting in experimental renal cell carcinoma models, and inhibition of fatty acid oxidation improved body weight and muscle mass in mice (Fukawa et al., 2016). While that study posited a central role for tumor cells, our data support adipose as a major source for the fatty acids metabolized in muscle in cachexia. Taken together, our results indicate that PDAC affects fat earlier and more profoundly than muscle, eliciting tissue inflammation and lipolysis, and that the consequent lipid accumulation in muscle alters muscle metabolism, leading to wasting. Consistent with this model, lipid accumulation in muscle portends poor outcomes in PDAC. Myosteatosis, sarcopenia and the combination are associated with reduced survival in one study of resectable patients with pancreatic and periampullary adenocarcinomas (Stretch et al., 2018), while only myosteatosis was associated with inflammation and reduced survival in another study of patients with unresectable pancreatic cancer or cholangiocarcinoma (Rollins et al., 2016). These results support clinical relevance of our observations here.

Our study also revealed that skeletal muscle talks back to adipose tissue in PDAC, apparently through production of soluble IL6R that accumulates in adipose tissue and activates STAT3. Soluble IL6R is largely produced by proteolytic cleavage of membrane GP80, which in mice and humans is mediated by ADAM10 and ADAM17 proteases (Schumacher et al., 2015). Muscle steatosis is correlated with increases in diacylglycerol (DAG), insulin resistance and increased activation of PKC theta (Szendroedi et al., 2014). Furthermore, PKC activation is a known rate limiting step for shedding of IL-6R (Mullberg et al., 1992). These previous findings suggest a connection between adipose wasting and induction of trans-signaling via muscle-derived sIL6R.

Finally, we propose that both steatosis and cytokines associated with inflammation augment muscle wasting. We tested our hypothesis by using exogenous mouse IL-6 protein and the fatty acid palmitate in excess to treat C2C12 myotubes. Indeed, treating myotubes with either IL-6 or palmitate alone induced significant myotube atrophy. However, the combination produced the most severe wasting between treatments. Taken together, our data suggest that cytokines and fatty acids conspire to effect muscle loss in PDAC.

Cancer cachexia is a systemic syndrome and understanding changes to tissue crosstalk in the presence of a tumor is critical for designing effective treatment. The data provided here strongly support targeted inhibition of IL-6R to treat PDAC cachexia. Pre-clinical studies document improved response to chemotherapy in mice with pancreatic cancer treated with anti-IL6R antibodies (Long et al., 2017); whether those mice also demonstrated reduced cachexia was not reported. An ongoing trial of the IL6R inhibitor Tocilizumab with gemcitabine and nab-paclitaxel in patients with advanced and metastatic disease might provide important insights on this approach in PDAC (Chen et al., 2017). As well, our results add to the literature implicating fat loss as a potential driver of cachexia mortality and immunomodulation in cancer (Tsoli et al., 2016), supporting the need to monitor adipose in patients with cachexia-associated cancers (Ebadi and Mazurak, 2015). Moreover, our studies also suggest that strategies as diverse as targeting tumor-cell production of IL-6, IL6R shedding from muscle and other tissues, adipose lipolysis, or lipid uptake by muscle might each show benefit in PDAC.

## ACKNOWLEDGMENTS

Images from Servier Medical Art in the Summary Figure are used under a Creative Commons license. This work was funded in part by grants to TAZ from the National Institutes of Health (Grants R01CA122596 and R01CA194593), the VA (Grant I01BX004177), the IU Simon Cancer Center, and the Lustgarten Foundation; to LGK from NIH (Grant R01DK096167) and the Lilly Endowment, Inc.; and to AB from NIH (Grant R21CA190028). Bioinformatics analysis was supported by IU Simon Cancer Center (Grant P30CA082709), Purdue University Center for Cancer Research (Grant P30CA023168) and Walther Cancer Foundation. JER was funded in part through an NIH training grant to Hal Broxmeyer (Grant T32 HL007910). We thank the Indiana University Simon Cancer Center for use of the Flow Cytometry Core, the Genomics Core, and Cancer Bioinformatics. Antibodies to myosin heavy chain and dystrophin were obtained from the Developmental Studies Hybridoma Bank, created by the NICHD of the NIH and maintained at The University of Iowa, Department of Biology, Iowa City, IA.

## AUTHOR CONTRIBUTIONS AND DECLARATIONS OF INTEREST

JER, AB, AN and YL carried out experiments. JER, LGK, and TAZ designed the studies. JER and TAZ wrote the manuscript. JER, AB, AN, TMO, LGK, and TAZ edited the manuscript.

The authors have no conflicts of interest to declare.

## Supplementary Figures

**Figure S1.**
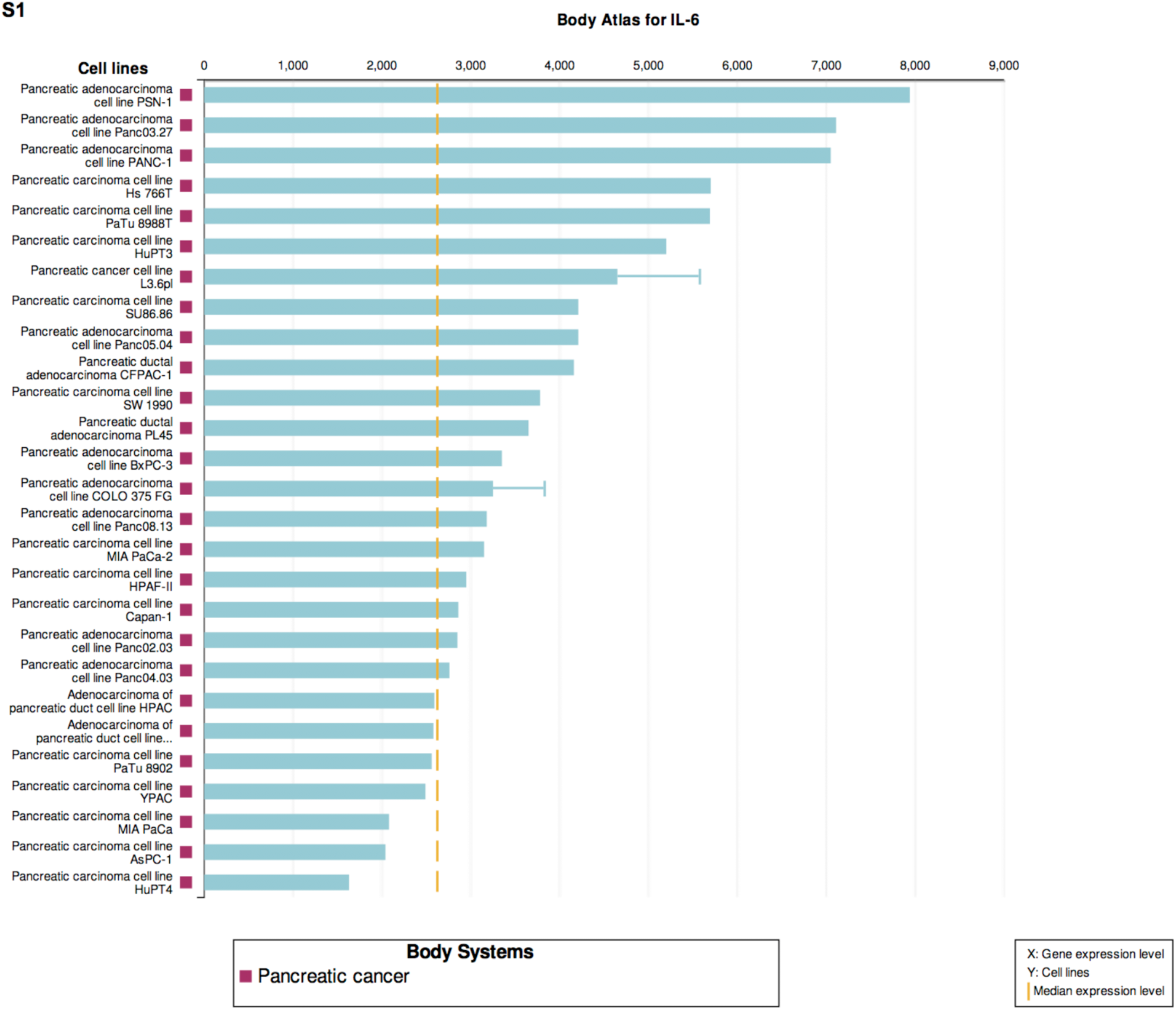
Expression of IL6 RNA in human pancreatic cancer cell lines. Expression levels of *IL6* mRNA across multiple human pancreatic cancer cell lines from Illumina BaseSpace Correlation Engine illustrating heterogeneity in *IL6* expression.

**Figure S2.**
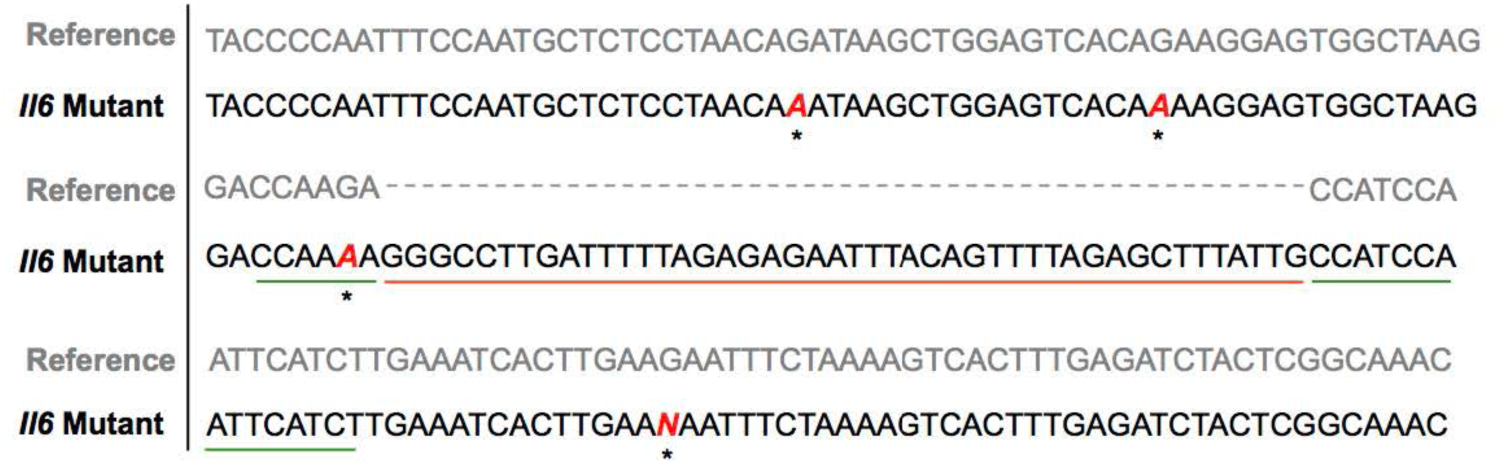
Characterization of KPC-IL-6^KO^ cells. DNA sequencing was performed at the CRISPR/Cas9 target site (green line) within the murine *Il6* gene showing an insertion mutation of 45 nucleotides (red line) into the KPC IL-6^KO^ mutant sequence. Asterisks indicate point mutations.

**Figure S3.**
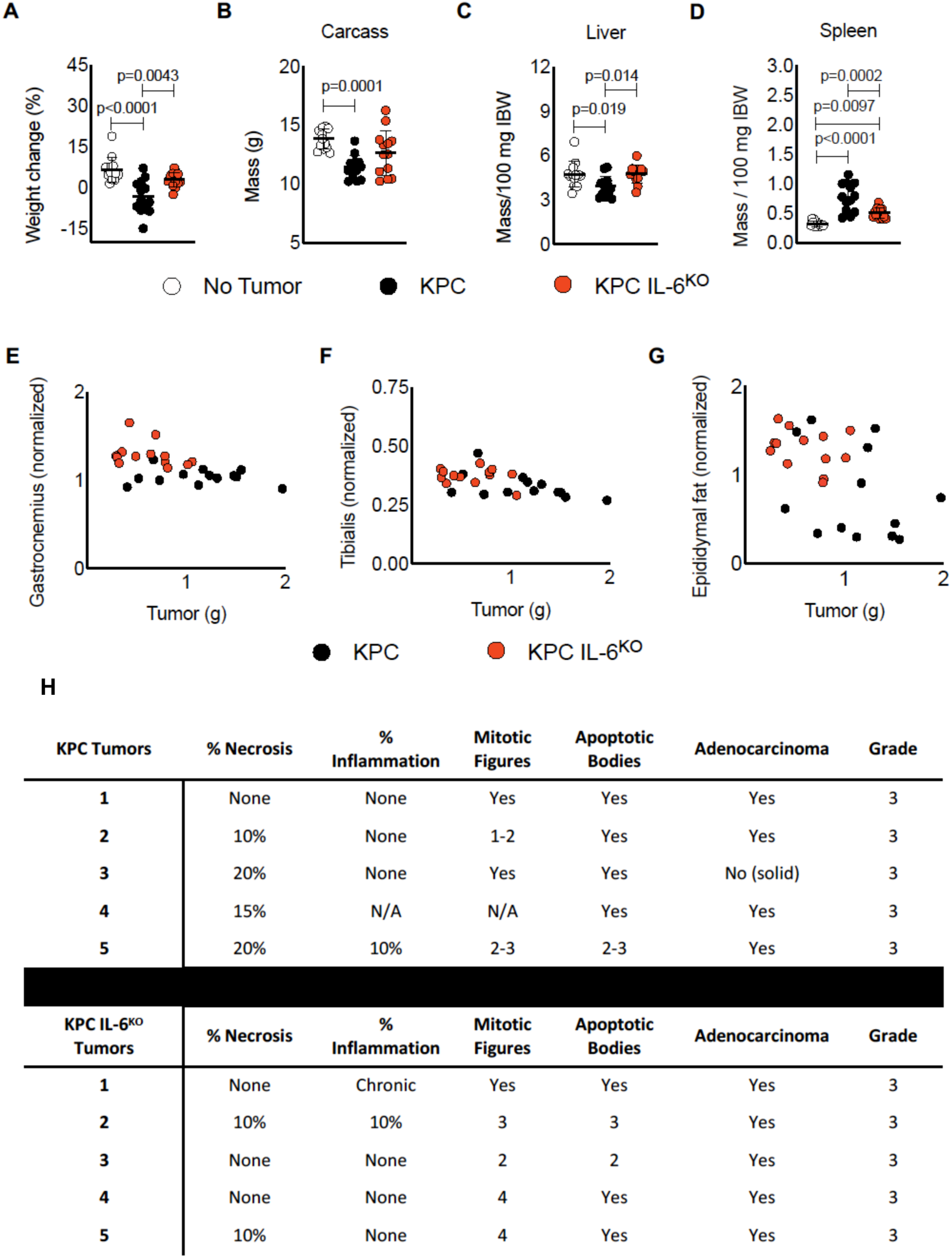
Deletion of tumor cell-derived IL-6 is associated with reduced weight loss, carcass and liver wasting, and splenomegaly in mice with PDAC. Tumor free final body weight was measured at euthanasia (**A**) and tissue weights for the carcass (**B**), liver (**C**) and spleen (**D**) were also recorded. No correlation was observed between tumor mass and gastrocnemius (**E**), tibialis (**F**), and epididymal fat pad (**G**) masses. Sections of excised KPC and KPC IL-6^KO^ tumors were stained with H&E and blindly scored by two separate pathologists to determine any differences in tumor grade (**H**).

## EXPERIMENTAL MODELS AND SUBJECT DETAILS

### Immunohistochemistry (IHC) performed on human and mouse tumors

Formalin-fixed human pancreatic ductal adenocarcinoma (PDAC) tumor arrays were obtained from US Biomax (PA1001a, Rockville, MD, USA) and consisted of 100 core samples from 50 patients. Twenty-eight core samples consisting of pancreas tissue adjacent to tumor tissue or tumor samples having no reactivity for IL-6 were excluded. Removal of these specimens was justified since adjacent tissue is not actually healthy, normal pancreas but rather tissue with a large degree of inflammation and no reactivity for IL-6 in either stromal or epithelial compartments is typically an indication of poor tissue procurement. The final sample population was made up of 72 core samples and included 21 males and 15 females ranging in age from 40-78 years of age. Tumor samples varied in stage from IB (maximum tumor diameter greater than four centimeters) to IV (metastases in four or more regional lymphnodes) as defined by the 8^th^ edition of AJCC cancer staging manual. Sections were deparaffinized with xylenes and rehydrated using deionized water (dH_2_O). Antigen retrieval was performed by incubating the sections in a sodium citrate buffer (10 mM sodium citrate, 0.05% Tween 20, pH 6.0) at 95° C for 10 minutes and allowing them to cool to room temperature for 30 minutes. Sections were washed using dH_2_O and endogenous peroxidase was quenched using 3% hydrogen peroxide (S25360, Thermo Fisher; Waltham, MA, USA). Sections incubated with blocking buffer which was comprised of 8% FBS (SH3007103HI, Thermo Fisher; Waltham, MA USA) in PBS (21040CV, Corning; Corning, NY, USA) for one hour and incubated overnight in a humid chamber at 4° C with an anti-IL-6 monoclonal antibody (mAb) (ab6672, Abcam; Cambridge, MA, USA) diluted 1:400 in blocking buffer. Detection and visualization of the anti-IL-6 mAb was done using ImmPACT DAB Peroxidase (HRP) Substrate kit from Vector Laboratories (SK-4105, Burlingame, CA, USA). Sections were counter stained with hematoxylin for one minute, dehydrated and cleared using sequential incubations (3 minutes each) of dH_2_O, 70% ethanol, 95% ethanol, 100% xylenes and then cover slips applied.

### In Vitro Cell Culture

C2C12 mouse myoblasts were purchased from ATCC (CRL-1772; Manassas, VA, USA). 3T3 fibroblasts were purchased from Zen Bio (SP-LF-1, Zen Bio; Research Triangle Park, NC, USA). KPC tumor cells were a gift from Dr. David Tuveson, who isolated them from a pancreatic tumor from a male LSL-**K**rasG12D: LSL-Tr**p**53R172H:Pdx1-**C**re (KPC) mouse on a C57BL/6 background (Hingorani et al., 2005) and we used these cells to generate the KPC IL-6^KO^ cells (RRID: CVCL_UR56).

#### i) C2C12 Mouse Myoblast Cell culture and Differentiation into Myotubes

Frozen murine C2C12 myoblasts (2 × 10^6^ cells) were thawed in a 37°C water bath and then seeded in flasks (130191, Thermo Scientific; Waltham, MA, USA) containing 20 mL of growth media (GM), which consisted of DMEM base media (10013CV, Corning; Corning, NY, USA), 10% fetal bovine serum (FBS) (SH30071.03, GE Healthcare Life Sciences; Pittsburgh, PA, USA) and 0.1% penicillin and streptomycin (15140122, ThermoFisher Scientific, Waltham, MA, USA;) and maintained in a humidified incubator at 37°C with 5% CO_2_ until 70% confluent (∼2 days). The GM was aspirated, the cells were washed with PBS and adherent cells dislodged using 3 mL of 0.25% trypsin (25053CL, Corning; Corning, NY, USA). Cells were then seeded 0.3 × 10^6^ cells per well in 6-well plates (140685, Thermo Scientific; Waltham, MA, USA) with growth media (GM) and maintained at 37°C with 5% CO_2_ until 90% confluent. The GM was then aspirated, cells were washed with PBS and 2 mL per well of differentiation media (DM) consisting of DMEM base media (10013CV, Corning; Corning, NY, USA), 2% horse serum (26050-088, Life Technologies; Carlsbad, CA, USA) and 0.1% penicillin and streptomycin (15140122, Gibco; Dublin, Ireland) was added and cells maintained at 37°C with 5% CO_2_. The DM was replaced every 48 hours and C2C12 myoblasts were fully differentiated into myotubes after 5-6 days in DM.

#### ii) 3T3 Fibroblast Cell Culture and Differentiation into 3T3 Adipocytes

Frozen 3T3 fibroblasts (0.5 × 10^6^) were thawed and seeded in flasks with preadipocyte media (PM-1-L1, Zen Bio; Research Triangle Park, NC, USA) at 37°C with 5% CO_2_ in a humidified incubator until 70% confluent. Adherent cells were then dislodged using 0.25% trypsin and seeded 0.3 × 10^6^ cells per well in six well plates with preadipocyte media and placed in the incubator. Cells were allowed to become 100% confluent, at which time cells were maintained at 100% confluence in preadipocyte media for an additional 48 hours to ensure growth arrest, refreshing the preadipocyte media every 48 hours. Cells were then washed with PBS and differentiation media (DM-2-L1, Zen Bio; Research Triangle Park, NC, USA) was added to the wells and cells placed back into the incubator. Cells were cultured for three days in differentiation media. Cells were then washed with PBS and adipocyte maintenance media (AM-1-L1, Zen Bio; Research Triangle Park, NC, USA) was added to the wells and cells placed back into the incubator. Cells were cultured in maintenance media for ten days refreshing the maintenance media every 48 hours. At ten days, fully differentiated 3T3 adipocytes were produced.

#### iii) Generation of KPC and KPC IL-6^KO^ Tumor Cell Lines

Frozen KPC cells were thawed and cultured flasks with GM as described for C2C12 myoblasts. When cells reached 70% confluence adherent cells were removed using trypsin (per C2C12 methods) and plated 0.3 × 10^6^ cells per well in two separate 6-well plates. Once cells were 70% confluent, Lipofectamine 3000 (L3000008, Invitrogen; Carlsbad, CA, USA) was used to transfect one plate of KPC cells with a CRISPR/Cas9 plasmid (CRISPR06-1EA, Sigma Aldrich; St. Louis, MO, USA) encoded with a green fluorescence protein (GFP) and null guide RNA. The additional plate was transfected with a CRISPR/Cas9 plasmid (MM0000468506, Sigma Aldrich; St. Louis, MO, USA) encoded with a GFP and a guide RNA specifically targeting the *mIl6* gene. Cells were incubated for 48 hours with the lipofectamine and CRISPR plasmids. Cells were removed from the wells using trypsin and sorted one GFP-positive cell per well into separate 96-well plates (167008, Thermo Fisher; Waltham, MA, USA) using fluorescence-activated cell sorting (FACS). One 96-well plate contained cells transfected with the null plasmid and the other plate contained cells transfected with the plasmid targeting *mIl6*. Single cell cultures were maintained in GM and expanded into colonies over ∼4 weeks. Colonies transfected with the plasmid targeting *mIl6* were screened for loss of *Il6* mRNA expression using RT-PCR. The selected clone with loss of *Il6* mRNA expression was designated KPC IL-6^KO^. Colonies transfected with the null plasmid were screened using RT-PCR for *Il6* mRNA expression similar to that of the parental cell line (untransfected) and the selected control clone was designated KPC.

#### iv) Proliferation Assay of KPC Cells

Comparison of *in vitro* proliferation between untransfected KPC cells (KPC-P), KPC cells transfected with a null CRISPR/Cas9 plasmid (KPC) and the mutant KPC IL-6^KO^ cells was performed using the xCELLigence Real-Time Cell Analysis (RTCA) assay (ACEA Biosciences; San Diego, CA). Cells were plated in GM at 2,000 cells per well in triplicate in an Eplate 16 (ACEA Biosciences; San Diego, CA). Cell growth was measured each hour for 100 hours.

### In Vivo Mouse Experiments

Experiments that included mice were approved by and performed in accordance with the Indiana University School of Medicine Institutional Animal Care and Use Committee. Seven-week-old male C57BL/6J mice (000664, Jackson Laboratory; Bar Harbor, Maine, USA) mice were group-housed in a barrier facility with ad libitum access to autoclaved food (Envigo: Huntingdon, Cambridgeshire, United Kingdom), sterile water, maintained on a 12hr light/dark cycle and allowed to acclimate to the facility for one week.

#### i) Orthotopic Implantation of KPC and KPC IL-6^KO^ Cells into Mice

Eight-week-old, male C57BL/6J mice were anesthetized using 4% isoflurane in an induction chamber within a biosafety cabinet. Mice were subcutaneously injected with buprenorphine (4202317005, PAR Pharmaceuticals; Chestnut Ridge, NY, USA) at 0.5 mg/kg in 0.1 mL sterile PBS prior to surgery and then every 12 hours from the first injection for 48 hours.

The abdomen was shaved, and mice were placed in lateral recumbency with the right side against a heated surgical platform and the left side facing upward. From this point forward aseptic technique was maintained. A sterile drape was placed over the abdomen and a hole cut in the drape to reveal the abdominal skin. The skin was prepped using three alternating scrubs of Betadine (6761815117, Purdue Products; Stamford, CT, USA) and 100% isopropanol. An incision (∼1 cm) was made through both skin and muscle into the left middle quadrant of the abdomen directly above the spleen using scissors. The pancreas was retracted using ring tip forceps by gripping the spleen (which is loosely connected with the pancreas) and retracting both the spleen and pancreas onto the sterile drape. The pancreas was injected with 5 × 10^4^ KPC or KPC IL-6^KO^ cells in 50μl GM of cells using a 1 mL syringe with a 26G needle (329652, Becton, Dickinson and Co.; Franklin Lakes, NJ, USA) over a period of thirty seconds. The spleen and accompanying pancreas were gently laid back into the abdominal cavity using the ring tip forceps. The incision was closed by suturing the abdominal wall musculature using absorbable 4.0-vicryl suture (33227, Butler Schein Animal Health; Dublin, OH, USA) and the skin flap closed using stainless steel wound clips (500346, World Precision Instruments; Sarasota, FL, USA).

## Experimental Details

### Euthanasia and Tissue Excision from Animal Models

All mice were euthanized when any group exhibited average body weight loss>10% or total fat mass <5% via EchoMRI. Euthanasia under general anesthesia (4% isoflurane) was done by exsanguination via cardiac puncture followed by cervical dislocation. Prior to euthanasia, blood was collected in EDTA tubes (366643, Becton, Dickinson and Co.; Franklin Lakes, NJ, USA) via cardiac puncture after anesthetizing the mice. The plasma was separated using centrifugation of the blood at 3,500 rpm at 4°C and stored at −80°C. Immediately following euthanasia tissues were excised and weighed. Sections of tissues were either placed into 2 mL cryogenic storage vials (82050-208, VWR; Radnor, PA, USA) and snap frozen in liquid nitrogen. Additionally, sections of tumor were placed into tubes containing 10% neutral buffered formalin (NBF) and sections of epididymal fat were placed into tubes containing Bouin’s solution 23-005-69, Thermo Fisher; Waltham, MA USA) for 24 hours. Fixed tumor and fat tissues were washed twice with PBS and transferred to new tubes containing 70% ethanol and stored at 4°C. Sections of the quadriceps muscle were mounted to cork discs, keeping the muscle fascicles perpendicular to the disc, by using Tissue plus O.C.T. compound (23-730-571, Thermo Healthcare; Waltham, MA USA). The muscle blocks were snap frozen by submerging the mounted muscle for 45 seconds in isopentane (2-methylbutane) (03551-4, Fisher Chemical; Waltham, MA USA) cooled to −150°C in a bath of liquid nitrogen and then stored at −80°C.

### RNA Isolation, Library Preparation, Sequencing and Quantitative PCR (qPCR)

RNA was isolated from ∼50 mg of snap frozen tissue or from cells cultured in 6- well plates via the miRNeasy Mini Kit (217004, Qiagen; Fredrick, MD USA). The concentration and quality of total RNA samples was first assessed using Agilent 2100 Bioanalyzer. A RIN (RNA Integrity Number) of six or higher was required to pass the quality control. Isolated RNA from quadriceps and epididymal fat pads was taken from four animals per group (i.e. No tumor, KPC tumor and KPC IL-6^KO^ tumor mice) and subjected to next generation sequencing. For adipose tissue, 500 nanograms (ng) of RNA per sample were used to prepare dual-indexed strand-specific cDNA library using TruSeq Stranded mRNA Library Prep Kit (Illumina; San Diego, CA, USA). For skeletal muscle, 100 ng of RNA per sample were used to prepare single-indexed strand specific cDNA library using TruSeq Nano DNA Library Prep kit.

The resulting libraries were assessed for quantity and size distribution using a Qubit and Agilent 2100 Bioanalyzer. 200 picomolar pooled RNA libraries were utilized per flow cell for clustering amplification on cBot using HiSeq 3000/4000 PE Cluster Kit and sequenced with 2×75bp paired-end configuration on HiSeq4000 (Illumina; San Diego, CA, USA) using the HiSeq 3000/4000 PE SBS Kit. A Phred quality score (Q score) was used to measure the quality of sequencing. More than 95% of the sequencing reads reached Q30 (99.9% base call accuracy) for both muscle and adipose tissues.

The sequencing data were first assessed using FastQC (Babraham Bioinformatics, Cambridge, UK) and then all sequenced libraries were mapped to the mm10 mouse genome using STAR RNA-seq aligner (Dobin et al., 2013). Uniquely mapped sequencing reads were assigned to mm10 UCSC reference genome. The data were normalized using the TMM (trimmed mean of M values) method. One muscle sample from the No Tumor group was identified as an outlier and removed from further analyses. Differential expression analysis was performed using edgeR (McCarthy et al., 2012; Robinson et al., 2010) and the false discovery rate (FDR) was computed from p-values using the Benjamini-Hochberg procedure. Differentially expressed genes were determined as having a fold change of ≥1.5 and an FDR of ≤ 0.05. Ingenuity Pathway Analysis (Qiagen; Fredrick, MD USA) was used for secondary analysis of RNA sequencing results. Data are deposited into GEO Datasets under series record GSE123310.

For qPCR analyses, 100 ng of isolated RNA was reverse transcribed into cDNA using the VERSO cDNA Synthesis Kit (AB1453B, Thermo Fisher; Waltham, MA USA). Taqman Universal Master Mix II (4440038, Thermo Fisher; Waltham, MA, USA) and gene specific Taqman probes (see oligonucleotides in resource table) were used to measure gene expression using cDNA in Light Cycler 480 96-well plates (04729692001, Roche; Indianapolis, IN, USA) on a Light Cycler 96 (Roche; Indianapolis, IN USA). For each sample, TATA binding protein (*Tbp)* gene expression was used to normalize the expression of the target gene. Each well contained one PCR reaction and reactions for each sample of cDNA were performed in triplicate for both target and control genes. Results from qPCR analyses were analyzed using the 2^-ΔΔCT^ method and reported as fold change.

### Western Blotting

Tissue protein lysates were made by homogenizing snap frozen tissue in ice cold RIPA buffer (Harlow and Lane, 2006) using a Polytron PT 10/35 homogenizer with PCU 11 controller (Kinematica; Luzern, Switzerland). Protein lysate concentration was measured using the Pierce^TM^ BCA Protein Assay Kit (PI23228, Thermo Scientific; Waltham, MA USA). Protein lysate was then added 1:1 to 2X sample buffer (125 mM Tris; pH 6.8, 4% (w/v) SDS, 20% glycerol, 100 mM DTT, 0.02% (w/v) bromophenol blue) and heated at 95^°^C for five minutes. Proteins were then separated via SDS-PAGE by loading 30 μg of protein from each sample into wells on a 4-15% Criterion^TM^ TGX^TM^ gel (5671084, Bio-Rad; Hercules, CA, USA) in running buffer (25 mM Tris, 192 mM glycine, 0.1% SDS) at 140 volts for one hour using a Power Pac HC (Bio-Rad; Hercules, CA, USA). The proteins were transferred to 0.2μm nitrocellulose membranes (1620233, Bio-Rad; Hercules, CA, USA) in ice-cold transfer buffer (25 mM Tris, 192 mM glycine, 20% (v/v) methanol, pH 8.3) at 100 volts for thirty minutes. Membranes were blocked using Sea Block (37527, Thermo Scientific; Waltham, MA USA) for one hour at room temperature on a shaker table. Proteins were detected using antigen specific primary antibodies (see antibodies in reagents table) diluted in Sea block with 0.1% Tween 20 (BP337-500, Fisher Scientific; Waltham, MA USA) and incubated with the membranes overnight at 4°C. Membranes were then washed twice with PBS and primary antibodies were visualized using florescent DyLight^TM^ secondary antibodies (Cell Signaling; Danvers, MA, USA) with specificity against the primary antibodies and imaged on an Odyssey CLx (LiCor; Lincoln, NE, USA). The quantification of target proteins was done by normalizing target protein expression to the loading control protein expression (α tubulin for muscle, β-actin for adipose tissue) specific to each membrane using Image Studio version 4.0 (LiCor; Lincoln, NE, USA). Normalized protein expression was then presented as fold-change versus no tumor bearing mice.

### Muscle Histology

For Oil Red O (ORO) staining, succinate dehydrogenase (SDH) reaction and IF performed on muscle, snap frozen mounted muscle blocks were serial sectioned (10 μm thickness) at −20^°^C on a Leica CM1860 cryostat (Buffalo Grove, IL, USA) and muscle cross-sections mounted to Superfrost Plus microscope slides (12-550-15, Thermo Scientific; Waltham, MA USA).

### Oil Red O Staining

ORO staining solution was prepared by dissolving 0.7 g of ORO (00625-25G, Sigma Aldrich; St. Louis, MO, USA) with 100 mL of 100% propylene glycol (398039, Sigma Aldrich; St. Louis, MO, USA) for five minutes at 100^°^C. The ORO staining was then maintained at 60^°^C until needed. Muscle cross-sections were post fixed in 10% NBF for 10 minutes at room temperature. Sections were washed for 15 minutes in running tap water, and then submerged in two separate bathes of 100% propylene glycol for five minutes each. Sections were then incubated in ORO staining solution for ten minutes with gentle shaking, washed in 85% propylene glycol for three minutes and rinsed using distilled water for three minutes. Sections were counterstained with Harris Modified Hematoxylin (SH26-500D, Fisher Chemical; Waltham, MA USA), rinsed with running tap water for ten minutes, rinsed with distilled water for three minutes and then counter slips were mounted using Prolong Gold Antifade Mountant (P36934, Life Technologies; Carlsbad, CA, USA). Oil Red O staining intensity was measured using established protocols (Mehlem et al., 2013).

### SDH Reaction

Glass Coplin jars were prewarmed to 37^°^C in an incubator prior to beginning the reaction. Succinic acid solution was made by adding 0.4 g of succinic acid (6106-21-4, Acros Organics; New Jersey, USA), 0.04 g nitroblue tetrazolium (N8129, Sigma Aldrich; St. Louis, MO, USA) and 0.001 g of phenazine methosulfate (P9625, Sigma Aldrich; St. Louis, MO, USA) to 40 mL of 0.1M Tris buffer and pH equilibrated to 7.01. The succinic acid solution was warmed to 37^°^C and the muscle sections were submerged in the solution for 30 minutes maintaining the temperature at 37^°^C in an incubator. Sections were then washed using a series of 30%, 60%, 90%, 60%, 30% acetone (A18-1, Fisher Chemical; Waltham, MA USA) solutions for 5 seconds each and rinsed in distilled water. Coverslips were applied using Prolong Gold Antifade mounting media. SDH intensity was measured using color brightfield on the Lionheart LX Automated Microscope (Biotek; Winooski, VT, USA).

### Measurement of C2C12 Myotube Diameter and Quadriceps Muscle Fiber Cross-sectional Area (CSA) Using Immunofluorescence (IF)

Fully differentiated myotubes were washed with PBS and fixed by incubation with ice-cold acetone and methanol (A412-4, Fisher Chemical; Waltham, MA USA) (1:1) at −20^°^C for 15 minutes. Myotubes were washed with PBS twice and blocked using blocking buffer (8% FBS in PBS) at room temperature for one hour. Myotubes were then incubated with an anti-myosin primary mAb (MF 20, DSHB; Iowa City, IA, USA) at a dilution of 1:50 in blocking buffer overnight at 4° C in a humid chamber. Myotubes were washed with PBS and incubated with a florescent secondary antibody (A-11029, Thermo Fisher; Waltham, MA USA) for one hour at room temperature in a humid chamber. Myotubes were washed with PBS, imaged and myotube diameter measured using Image J analysis.

IF was performed on frozen muscle cross-sections that were post-fixed in pre-cooled acetone (-20^°^C) for ten minutes and placed at room temperature for 30 minutes. Sections were washed twice in PBS and then blocked by incubating the sections with blocking buffer for one hour at room temperature. The sections were washed with PBS and incubated with the primary antibodies against IL6R and CD31 (see table for specific antibody details) diluted in blocking buffer overnight at 4^°^C in a humid chamber.

Sections were washed and the secondary antibody applied (see table for specific antibody details) at a dilution of 1:1000 in blocking buffer for one hour at room temperature. Sections were washed and incubated with DAPI (268298, EMD Millipore; Burlington, MA, USA) for 1 minute to visualize nuclei. Sections were washed and cover slips applied using Prolong Gold Antifade mounting media.

To measure muscle fiber CSA using IF, frozen sections of mouse quadriceps muscles were fixed and blocked as described previously and incubated with an anti-dystrophin primary antibody (MANDRA11, DSHB; Iowa City, IA, USA) and detection of the primary antibody was done using a goat anti-mouse AlexaFluor 594 florescent secondary antibody (A-11032, Thermo Fisher; Waltham, MA, USA). Muscle fiber CSA was measured from images taken of the IF dystrophin sections using an ImageJ macro obtained from the Lieber laboratory (Mehlem et al., 2013; Minamoto et al., 2007).

### Mouse Tumor and Adipose Tissue IHC

Tumors were excised from mice and immediately fixed in 10% neutral buffered formalin (NBF) for 48 hours. Tumors were then washed in PBS and stored in 70% ethanol at 4° C until analysis. Tumors were paraffin embedded and cross-sections were cut at 5 μm and mounted to microscope slides. Deparaffinization, antigen retrieval, endogenous peroxidase neutralization and blocking were performed as previously described for human tumors. After blocking, tumors were incubated with an anti-IL-6 mAb (ab7737, Abcam; Cambridge, MA, USA) at 1:50 dilution in blocking buffer at 4° C overnight in a humid chamber. Detection of the primary mAb, counter staining, clearing and cover slip application were performed as described for the human tumors.

Fixed adipose tissue was washed in PBS and stored in 70% ethanol at 4° C. Fixed adipose tissue was then paraffin embedded and sectioned at 5 μm and mounted to microscope slides. Deparaffinization, antigen retrieval, endogenous peroxidase neutralization and blocking were performed as previously described for human tumors. Sections were then incubated with an anti-IL-6 mAb (ab7737, Abcam; Cambridge, MA, USA) diluted 1:50 in blocking buffer or an anti-F4/80 (NB600-404, Novus Biologicals; Centennial, CO, USA) diluted 1:250 in blocking buffer overnight at 4° C in a humid chamber. Detection of the primary mAb, counter staining, clearing and cover slip application were performed as described for the human tumors.

### Plasma Analysis of IL-6, IL6R, Glycerol, and Fatty Acids

Mouse plasma obtained from cardiac puncture was measured for IL-6 and IL6R protein levels via enzyme-linked immunosorbent assay (ELISA) mouse Quantikine kits (MR600 & M6000B, R&D Systems; Minneapolis, MN USA). Plasma glycerol and FAs were measured using Glycerol and FA Quantitation Kits (MAK117 & MAK044, Sigma Aldrich; St. Louis, MO USA).

### Treatment of C2C12 Myotubes and 3T3 Adipocytes with KPC Conditioned Media (CM) and Measurement of Il6 and Il6ra mRNA

C2C12 myoblasts and 3T3 fibroblasts were cultured separately in six well plates and differentiated into myotubes and adipocytes, respectively (see *in vitro* methods). KPC cells were cultured in GM in flasks until becoming 100% confluent. At that time, the GM was refreshed and 100% confluent KPC cells were maintained for an additional 24 hours. After 24 hours, the CM was collected, centrifuged at 3,500 rpm to pellet unwanted cellular debris and the supernatant collected. Myotubes and adipocytes were treated in triplicate with either 30% GM (control) or 30% KPC CM per well. For myotubes, the treatment media was aspirated, cells were washed and RNA from myotubes was isolated at 12 hours, 48 hours and 96 hours after treatment. For 3T3 adipocytes, RNA was isolated at one hour and three hours post-treatment in the same method used for the myotubes. The timepoints selected for 3T3 adipocytes were selected from established protocols for the measurement of lipolysis (Schweiger et al., 2014). Myotube and adipocyte expression of *Il6* and *Il6r* mRNA at each timepoint was measured via qPCR using probes for *Il6* (Mm00446190, Applied Biosciences; Beverly Hills, CA, USA) and *Il6r* (Mm01211444, Applied Biosciences; Beverly Hills, CA, USA).

### Treatment of C2C12 myotubes with CM from 3T3 adipocytes treated with KPC CM and treatment of 3T3 adipocytes with CM from C2C12 myotubes treated with KPC CM

C2C12 myoblasts and 3T3 fibroblasts were cultured separately in six well plates and differentiated into myotubes and adipocytes, respectively (see *in vitro* methods). Mature adipocytes were treated with 30% KPC CM for one hour, the KPC CM was removed, adipocytes were washed with PBS and then normal adipocyte media was applied to collect the adipocyte secretome in response to KPC CM stimulation. This adipocyte CM was then harvested at one hour and three hours after application to KPC CM stimulated adipocytes. Myotubes were then treated with 30% adipocyte CM for 48 hours and myotube diameter measured.

Mature C2C12 myotubes were treated with 30% KPC CM for 12 hours, 48 hours and 96 hours and the myotube CM was collected at each of these time points. Mature 3T3 adipocytes were then treated with 30% C2C12 myotube CM from each timepoint (ie, 12, 48, 96 hours) for one hour, C2C12 CM was then removed, adipocytes were washed with PBS and normal adipocyte media was applied. Normal adipocyte media was harvested at one hour and three hours after application and measured for glycerol content as a marker for lipolysis.

### Treatment of Myotubes with IL-6, IL6R and Neutralizing Antibodies

C2C12 myoblasts were differentiated into myotubes in six well plates. Fully differentiated myotubes were then treated in triplicate with either DF media (control), recombinant mouse IL-6 (300 pg/mL) protein (406-ML-005, R&D Systems; Minneapolis, MN USA), recombinant mouse IL6R (25 ng/mL) (P9767-5G, Sigma Aldrich; St. Louis, MO, USA), the combination of IL-6 (300 pg/mL) and IL6R (25 ng/mL) (1830-SR-025, R&D Systems; Minneapolis, MN, USA) or IL-6 (300 pg/mL) in the presence of IL-6 neutralizing antibody (2 mg/mL) (14-7061-85, Thermo Fisher; Waltham, MA, USA) for 48 hours. After 48 hours of treatment, myotubes were washed with PBS and fixed using ice-cold acetone/methanol (1:1) for fifteen minutes at −20° C. Myotubes were then visualized with IF using an anti-myosin antibody (see IF methods) and myotube diameter measured with Image J.

### Statistical Analyses

The Student t-test was used for the comparison of means between two datasets while one-way ANOVA with the Tukey’s post-hoc test was used for comparison of the means between three or more datasets. The differences between group means was considered statistically different when P<0.05. Data are presented as mean ± SEM. All statistical analyses were performed using Graph Pad Prism version 7.0 (San Diego, CA, USA).

## REFERENCES CITED

Acharyya, S., M.E. Butchbach, Z. Sahenk, H. Wang, M. Saji, M. Carathers, M.D. Ringel, R.J. Skipworth, K.C. Fearon, M.A. Hollingsworth, P. Muscarella, A.H. Burghes, J.A. Rafael-Fortney, and D.C. Guttridge. 2005. Dystrophin glycoprotein complex dysfunction: a regulatory link between muscular dystrophy and cancer cachexia. Cancer Cell 8:421–432.

Argiles, J.M., B. Stemmler, F.J. Lopez-Soriano, and S. Busquets. 2018. Inter-tissue communication in cancer cachexia. Nat Rev Endocrinol 15:9–20.

Babic, A., N. Schnure, N.P. Neupane, M.M. Zaman, N. Rifai, M.W. Welch, L.K. Brais, D.A. Rubinson, V. Morales-Oyarvide, C. Yuan, S. Zhang, E.M. Poole, B.M. Wolpin, M.H. Kulke, D.A. Barbie, K. Wong, C.S. Fuchs, and K. Ng. 2018. Plasma inflammatory cytokines and survival of pancreatic cancer patients. Clin Transl Gastroenterol 9:145.

Baltgalvis, K.A., F.G. Berger, M.M. Pena, J.M. Davis, S.J. Muga, and J.A. Carson. 2008. Interleukin-6 and cachexia in ApcMin/+ mice. Am J Physiol Regul Integr Comp Physiol 294:R393–401.

Baltgalvis, K.A., F.G. Berger, M.M. Pena, J.M. Davis, J.P. White, and J.A. Carson. 2009. Muscle wasting and interleukin-6-induced atrogin-I expression in the cachectic Apc (Min/+) mouse. Pflugers Arch 457:989–1001.

Begue, G., A. Douillard, O. Galbes, B. Rossano, B. Vernus, R. Candau, and G. Py. 2013. Early activation of rat skeletal muscle IL-6/STAT1/STAT3 dependent gene expression in resistance exercise linked to hypertrophy. PLoS One 8:e57141.

Bohnert, K.R., Y.S. Gallot, S. Sato, G. Xiong, S.M. Hindi, and A. Kumar. 2016. Inhibition of ER stress and unfolding protein response pathways causes skeletal muscle wasting during cancer cachexia. FASEB J 30:3053–3068.

Bonetto, A., T. Aydogdu, X. Jin, Z. Zhang, R. Zhan, L. Puzis, L.G. Koniaris, and T.A. Zimmers. 2012. JAK/STAT3 pathway inhibition blocks skeletal muscle wasting downstream of IL-6 and in experimental cancer cachexia. Am J Physiol Endocrinol Metab 303:E410–421.

Bonetto, A., T. Aydogdu, N. Kunzevitzky, D.C. Guttridge, S. Khuri, L.G. Koniaris, and T.A. Zimmers. 2011. STAT3 activation in skeletal muscle links muscle wasting and the acute phase response in cancer cachexia. PLoS One 6:e22538.

Camargo, R.G., D.M. Riccardi, H.Q. Ribeiro, L.C. Carnevali, Jr., E.M. de Matos-Neto, L. Enjiu, R.X. Neves, J.D. Lima, R.G. Figueredo, P.S. de Alcantara, L. Maximiano, J. Otoch, M. Batista, Jr., G. Puschel, and M. Seelaender. 2015. NF-kappaBp65 and Expression of Its Pro-Inflammatory Target Genes Are Upregulated in the Subcutaneous Adipose Tissue of Cachectic Cancer Patients. Nutrients 7:4465–4479.

Camus, V., H. Lanic, J. Kraut, R. Modzelewski, F. Clatot, J.M. Picquenot, N. Contentin, P. Lenain, L. Groza, E. Lemasle, C. Fronville, N. Cardinael, M.L. Fontoura, A. Chamseddine, O. Brehar, A. Stamatoullas, S. Lepretre, H. Tilly, and F. Jardin. 2014. Prognostic impact of fat tissue loss and cachexia assessed by computed tomography scan in elderly patients with diffuse large B-cell lymphoma treated with immunochemotherapy. Eur J Haematol 93:9–18.

Chen, I., J.S. Johansen, T.A. Zimmers, C. Dehlendorff, V. Kirk Parner, B. Vittrup Jensen, and D. Nielsen. 2017. PACTO: A single center, randomized, phase II study of the combination of nab-paclitaxel and gemcitabine with or without tocilizumab, an IL-6R inhibitor, as first-line treatment in patients with locally advanced or metastatic pancreatic cancer Ann Oncol 28:mdx369.158.

Chen, J.L., K.L. Walton, H. Qian, T.D. Colgan, A. Hagg, M.J. Watt, C.A. Harrison, and P. Gregorevic. 2016. Differential Effects of IL6 and Activin A in the Development of Cancer-Associated Cachexia. Cancer Res 76:5372–5382.

Dalal, S., D. Hui, L. Bidaut, K. Lem, E. Del Fabbro, C. Crane, C.C. Reyes-Gibby, D. Bedi, and E. Bruera. 2012. Relationships among body mass index, longitudinal body composition alterations, and survival in patients with locally advanced pancreatic cancer receiving chemoradiation: a pilot study. J Pain Symptom Manage 44:181–191.

Danai, L.V., A. Babic, M.H. Rosenthal, E.A. Dennstedt, A. Muir, E.C. Lien, J.R. Mayers, K. Tai, A.N. Lau, P. Jones-Sali, C.M. Prado, G.M. Petersen, N. Takahashi, M. Sugimoto, J.J. Yeh, N. Lopez, N. Bardeesy, C. Fernandez-Del Castillo, A.S. Liss, A.C. Koong, J. Bui, C. Yuan, M.W. Welch, L.K. Brais, M.H. Kulke, C. Dennis, C.B. Clish, B.M. Wolpin, and M.G. Vander Heiden. 2018. Altered exocrine function can drive adipose wasting in early pancreatic cancer. Nature 558:600–604.

Das, S.K., S. Eder, S. Schauer, C. Diwoky, H. Temmel, B. Guertl, G. Gorkiewicz, K.P. Tamilarasan, P. Kumari, M. Trauner, R. Zimmermann, P. Vesely, G. Haemmerle, R. Zechner, and G. Hoefler. 2011. Adipose triglyceride lipase contributes to cancer-associated cachexia. Science 333:233–238.

Di Sebastiano, K.M., L. Yang, K. Zbuk, R.K. Wong, T. Chow, D. Koff, G.R. Moran, and M. Mourtzakis. 2013. Accelerated muscle and adipose tissue loss may predict survival in pancreatic cancer patients: the relationship with diabetes and anaemia. Br J Nutr 109:302–312.

Dobin, A., C.A. Davis, F. Schlesinger, J. Drenkow, C. Zaleski, S. Jha, P. Batut, M. Chaisson, and T.R. Gingeras. 2013. STAR: ultrafast universal RNA-seq aligner. Bioinformatics 29:15–21.

Ebadi, M., and V.C. Mazurak. 2015. Potential Biomarkers of Fat Loss as a Feature of Cancer Cachexia. Mediators Inflamm 2015:820934.

Ebrahimi, B., S.L. Tucker, D. Li, J.L. Abbruzzese, and R. Kurzrock. 2004. Cytokines in pancreatic carcinoma: correlation with phenotypic characteristics and prognosis. Cancer 101:2727–2736.

Flint, T.R., T. Janowitz, C.M. Connell, E.W. Roberts, A.E. Denton, A.P. Coll, D.I. Jodrell, and D.T. Fearon. 2016. Tumor-Induced IL-6 Reprograms Host Metabolism to Suppress Anti-tumor Immunity. Cell Metab 24:672–684.

Fukawa, T., B.C. Yan-Jiang, J.C. Min-Wen, E.T. Jun-Hao, D. Huang, C.N. Qian, P. Ong, Z. Li, S. Chen, S.Y. Mak, W.J. Lim, H.O. Kanayama, R.E. Mohan, R.R. Wang, J.H. Lai, C. Chua, H.S. Ong, K.K. Tan, Y.S. Ho, I.B. Tan, B.T. Teh, and N. Shyh-Chang. 2016. Excessive fatty acid oxidation induces muscle atrophy in cancer cachexia. Nat Med 22:666–671.

Harlow, E., and D. Lane. 2006. Lysing tissue-culture cells for immunoprecipitation. CSH Protoc 2006:

Hendifar, A.E., M.Q.B. Petzel, T.A. Zimmers, C.S. Denlinger, L.M. Matrisian, V.J. Picozzi, L. Rahib, and C. Precision Promise. 2018. Pancreas Cancer-Associated Weight Loss. Oncologist

Hingorani, S.R., L. Wang, A.S. Multani, C. Combs, T.B. Deramaudt, R.H. Hruban, A.K. Rustgi, S. Chang, and D.A. Tuveson. 2005. Trp53R172H and KrasG12D cooperate to promote chromosomal instability and widely metastatic pancreatic ductal adenocarcinoma in mice. Cancer Cell 7:469–483.

Holmer, R., F.A. Goumas, G.H. Waetzig, S. Rose-John, and H. Kalthoff. 2014. Interleukin-6: a villain in the drama of pancreatic cancer development and progression. Hepatobiliary Pancreat Dis Int 13:371–380.

Jin, X., T.A. Zimmers, E.A. Perez, R.H. Pierce, Z. Zhang, and L.G. Koniaris. 2006. Paradoxical effects of short- and long-term interleukin-6 exposure on liver injury and repair. Hepatology 43:474–484.

Kays, J.K., S. Shahda, M. Stanley, T.M. Bell, B.H. O’Neill, M.D. Kohli, M.E. Couch, L.G. Koniaris, and T.A. Zimmers. 2018. Three cachexia phenotypes and the impact of fat-only loss on survival in FOLFIRINOX therapy for pancreatic cancer. J Cachexia Sarcopenia Muscle 9:673–684.

Koniaris, L.G., I.H. McKillop, S.I. Schwartz, and T.A. Zimmers. 2003. Liver regeneration. J Am Coll Surg 197:634–659.

Kraakman, M.J., H.L. Kammoun, T.L. Allen, V. Deswaerte, D.C. Henstridge, E. Estevez, V.B. Matthews, B. Neill, D.A. White, A.J. Murphy, L. Peijs, C. Yang, S. Risis, C.R. Bruce, X.J. Du, A. Bobik, R.S. Lee-Young, B.A. Kingwell, A. Vasanthakumar, W. Shi, A. Kallies, G.I. Lancaster, S. Rose-John, and M.A. Febbraio. 2015. Blocking IL-6 trans-signaling prevents high-fat diet-induced adipose tissue macrophage recruitment but does not improve insulin resistance. Cell Metab 21:403–416.

Lesina, M., M.U. Kurkowski, K. Ludes, S. Rose-John, M. Treiber, G. Kloppel, A. Yoshimura, W. Reindl, B. Sipos, S. Akira, R.M. Schmid, and H. Algul. 2011. Stat3/Socs3 activation by IL-6 transsignaling promotes progression of pancreatic intraepithelial neoplasia and development of pancreatic cancer. Cancer Cell 19:456–469.

Long, K.B., G. Tooker, E. Tooker, S.L. Luque, J.W. Lee, X. Pan, and G.L. Beatty. 2017. IL6 Receptor Blockade Enhances Chemotherapy Efficacy in Pancreatic Ductal Adenocarcinoma. Mol Cancer Ther 16:1898–1908.

Mace, T.A., R. Shakya, J.R. Pitarresi, B. Swanson, C.W. McQuinn, S. Loftus, E. Nordquist, Z. Cruz-Monserrate, L. Yu, G. Young, X. Zhong, T.A. Zimmers, M.C. Ostrowski, T. Ludwig, M. Bloomston, T. Bekaii-Saab, and G.B. Lesinski. 2018. IL-6 and PD-L1 antibody blockade combination therapy reduces tumour progression in murine models of pancreatic cancer. Gut 67:320–332.

Martignoni, M.E., P. Kunze, W. Hildebrandt, B. Kunzli, P. Berberat, T. Giese, O. Kloters, J. Hammer, M.W. Buchler, N.A. Giese, and H. Friess. 2005. Role of mononuclear cells and inflammatory cytokines in pancreatic cancer-related cachexia. Clin Cancer Res 11:5802–5808.

Martin, L., L. Birdsell, N. Macdonald, T. Reiman, M.T. Clandinin, L.J. McCargar, R. Murphy, S. Ghosh, M.B. Sawyer, and V.E. Baracos. 2013. Cancer cachexia in the age of obesity: skeletal muscle depletion is a powerful prognostic factor, independent of body mass index. J Clin Oncol 31:1539–1547.

McCarthy, D.J., Y. Chen, and G.K. Smyth. 2012. Differential expression analysis of multifactor RNA-Seq experiments with respect to biological variation. Nucleic Acids Res 40:4288–4297.

McKay, B.R., M. De Lisio, A.P. Johnston, C.E. O’Reilly, S.M. Phillips, M.A. Tarnopolsky, and G. Parise. 2009. Association of interleukin-6 signalling with the muscle stem cell response following muscle-lengthening contractions in humans. PLoS One 4:e6027.

Mehlem, A., C.E. Hagberg, L. Muhl, U. Eriksson, and A. Falkevall. 2013. Imaging of neutral lipids by oil red O for analyzing the metabolic status in health and disease. Nat Protoc 8:1149–1154.

Melenovsky, V., M. Kotrc, B.A. Borlaug, T. Marek, J. Kovar, I. Malek, and J. Kautzner. 2013. Relationships between right ventricular function, body composition, and prognosis in advanced heart failure. J Am Coll Cardiol 62:1660–1670.

Minamoto, V.B., J.B. Hulst, M. Lim, W.J. Peace, S.N. Bremner, S.R. Ward, and R.L. Lieber. 2007. Increased efficacy and decreased systemic-effects of botulinum toxin A injection after active or passive muscle manipulation. Dev Med Child Neurol 49:907–914.

Moses, A.G., J. Maingay, K. Sangster, K.C. Fearon, and J.A. Ross. 2009. Pro-inflammatory cytokine release by peripheral blood mononuclear cells from patients with advanced pancreatic cancer: relationship to acute phase response and survival. Oncol Rep 21:1091–1095.

Mullberg, J., H. Schooltink, T. Stoyan, P.C. Heinrich, and S. Rose-John. 1992. Protein kinase C activity is rate limiting for shedding of the interleukin-6 receptor. Biochem Biophys Res Commun 189:794–800.

Narsale, A.A., and J.A. Carson. 2014. Role of interleukin-6 in cachexia: therapeutic implications. Curr Opin Support Palliat Care 8:321–327.

Ohlund, D., A. Handly-Santana, G. Biffi, E. Elyada, A.S. Almeida, M. Ponz-Sarvise, V. Corbo, T.E. Oni, S.A. Hearn, E.J. Lee, Chio, II, C.I. Hwang, H. Tiriac, L.A. Baker, D.D. Engle, C. Feig, A. Kultti, M. Egeblad, D.T. Fearon, J.M. Crawford, H. Clevers, Y. Park, and D.A. Tuveson. 2017. Distinct populations of inflammatory fibroblasts and myofibroblasts in pancreatic cancer. J Exp Med 214:579–596.

Okada, S., T. Okusaka, H. Ishii, A. Kyogoku, M. Yoshimori, N. Kajimura, K. Yamaguchi, and T. Kakizoe. 1998. Elevated serum interleukin-6 levels in patients with pancreatic cancer. Jpn J Clin Oncol 28:12–15.

Onesti, J.K., and D.C. Guttridge. 2014. Inflammation based regulation of cancer cachexia. Biomed Res Int 2014:168407.

Pedroso, F.E., P.B. Spalding, M.C. Cheung, R. Yang, J.C. Gutierrez, A. Bonetto, R. Zhan, H.L. Chan, N. Namias, L.G. Koniaris, and T.A. Zimmers. 2012. Inflammation, organomegaly, and muscle wasting despite hyperphagia in a mouse model of burn cachexia. J Cachexia Sarcopenia Muscle 3:199–211.

Ramsey, M.L., E. Talbert, D. Ahn, T. Bekaii-Saab, N. Badi, P.M. Bloomston, D.L. Conwell, Z. Cruz-Monserrate, M. Dillhoff, M.R. Farren, A. Hinton, S.G. Krishna, G.B. Lesinski, T. Mace, A. Manilchuk, A. Noonan, T.M. Pawlik, P.V. Rajasekera, C. Schmidt, D. Guttridge, and P.A. Hart. 2019. Circulating interleukin-6 is associated with disease progression, but not cachexia in pancreatic cancer. Pancreatology 19:80–87.

Razidlo, G.L., K.M. Burton, and M.A. McNiven. 2018. Interleukin-6 promotes pancreatic cancer cell migration by rapidly activating the small GTPase CDC42. J Biol Chem 293:11143–11153.

Robinson, M.D., D.J. McCarthy, and G.K. Smyth. 2010. edgeR: a Bioconductor package for differential expression analysis of digital gene expression data. Bioinformatics 26:139–140.

Rohm, M., M. Schafer, V. Laurent, B.E. Ustunel, K. Niopek, C. Algire, O. Hautzinger, T.P. Sijmonsma, A. Zota, D. Medrikova, N.S. Pellegata, M. Ryden, A. Kulyte, I. Dahlman, P. Arner, N. Petrovic, B. Cannon, E.Z. Amri, B.E. Kemp, G.R. Steinberg, P. Janovska, J. Kopecky, C. Wolfrum, M. Bluher, M. Berriel Diaz, and S. Herzig. 2016. An AMP-activated protein kinase-stabilizing peptide ameliorates adipose tissue wasting in cancer cachexia in mice. Nat Med 22:1120–1130.

Rollins, K.E., N. Tewari, A. Ackner, A. Awwad, S. Madhusudan, I.A. Macdonald, K.C. Fearon, and D.N. Lobo. 2016. The impact of sarcopenia and myosteatosis on outcomes of unresectable pancreatic cancer or distal cholangiocarcinoma. Clin Nutr 35:1103–1109.

Rosean, T.R., V.S. Tompkins, G. Tricot, C.J. Holman, A.K. Olivier, F. Zhan, and S. Janz. 2014. Preclinical validation of interleukin 6 as a therapeutic target in multiple myeloma. Immunol Res 59:188–202.

Sandini, M., M. Patino, C.R. Ferrone, C.A. Alvarez-Perez, K.C. Honselmann, S. Paiella, M. Catania, L. Riva, G. Tedesco, R. Casolino, A. Auriemma, M.C. Salandini, G. Carrara, G. Cristel, A. Damascelli, D. Ippolito, M. D’Onofrio, K.D. Lillemoe, C. Bassi, M. Braga, L. Gianotti, D. Sahani, and C. Fernandez-Del Castillo. 2018. Association Between Changes in Body Composition and Neoadjuvant Treatment for Pancreatic Cancer. JAMA Surg 153:809–815.

Sandri, M. 2016. Protein breakdown in cancer cachexia. Semin Cell Dev Biol 54:11–19.

Schaper, F., and S. Rose-John. 2015. Interleukin-6: Biology, signaling and strategies of blockade. Cytokine Growth Factor Rev 26:475–487.

Schumacher, N., D. Meyer, A. Mauermann, J. von der Heyde, J. Wolf, J. Schwarz, K. Knittler, G. Murphy, M. Michalek, C. Garbers, J.W. Bartsch, S. Guo, B. Schacher, P. Eickholz, A. Chalaris, S. Rose-John, and B. Rabe. 2015. Shedding of Endogenous Interleukin-6 Receptor (IL-6R) Is Governed by A Disintegrin and Metalloproteinase (ADAM) Proteases while a Full-length IL-6R Isoform Localizes to Circulating Microvesicles. J Biol Chem 290:26059–26071.

Schweiger, M., T.O. Eichmann, U. Taschler, R. Zimmermann, R. Zechner, and A. Lass. 2014. Measurement of lipolysis. Methods Enzymol 538:171–193.

Siegel, R.L., K.D. Miller, and A. Jemal. 2019. Cancer statistics, 2019. CA Cancer J Clin 69:7–34.

Stephens, N.A., R.J. Skipworth, I.J. Gallagher, C.A. Greig, D.C. Guttridge, J.A. Ross, and K.C. Fearon. 2015. Evaluating potential biomarkers of cachexia and survival in skeletal muscle of upper gastrointestinal cancer patients. J Cachexia Sarcopenia Muscle 6:53–61.

Stretch, C., J.M. Aubin, B. Mickiewicz, D. Leugner, T. Al-Manasra, E. Tobola, S. Salazar, F.R. Sutherland, C.G. Ball, E. Dixon, H.J. Vogel, S. Damaraju, V.E. Baracos, and O.F. Bathe. 2018. Sarcopenia and myosteatosis are accompanied by distinct biological profiles in patients with pancreatic and periampullary adenocarcinomas. PLoS One 13:e0196235.

Suh, S.Y., Y.S. Choi, C.H. Yeom, S.M. Kwak, H.M. Yoon, D.G. Kim, S.J. Koh, J. Park, M.A. Lee, Y.J. Lee, A.R. Seo, H.Y. Ahn, and E. Yim. 2013. Interleukin-6 but not tumour necrosis factor-alpha predicts survival in patients with advanced cancer. Support Care Cancer 21:3071–3077.

Sun, L., X.Q. Quan, and S. Yu. 2015. An Epidemiological Survey of Cachexia in Advanced Cancer Patients and Analysis on Its Diagnostic and Treatment Status. Nutr Cancer 67:1056–1062.

Szendroedi, J., T. Yoshimura, E. Phielix, C. Koliaki, M. Marcucci, D. Zhang, T. Jelenik, J. Muller, C. Herder, P. Nowotny, G.I. Shulman, and M. Roden. 2014. Role of diacylglycerol activation of PKCtheta in lipid-induced muscle insulin resistance in humans. Proc Natl Acad Sci U S A 111:9597–9602.

Talbert, E.E., H.L. Lewis, M.R. Farren, M.L. Ramsey, J.M. Chakedis, P. Rajasekera, E. Haverick, A. Sarna, M. Bloomston, T.M. Pawlik, T.A. Zimmers, G.B. Lesinski, P.A. Hart, M.E. Dillhoff, C.R. Schmidt, and D.C. Guttridge. 2018. Circulating monocyte chemoattractant protein-1 (MCP-1) is associated with cachexia in treatment-naive pancreatic cancer patients. J Cachexia Sarcopenia Muscle 9:358–368.

Taniguchi, K., and M. Karin. 2014. IL-6 and related cytokines as the critical lynchpins between inflammation and cancer. Semin Immunol 26:54–74.

Trujillo, M.E., S. Sullivan, I. Harten, S.H. Schneider, A.S. Greenberg, and S.K. Fried. 2004. Interleukin-6 regulates human adipose tissue lipid metabolism and leptin production in vitro. J Clin Endocrinol Metab 89:5577–5582.

Tsoli, M., M.M. Swarbrick, and G.R. Robertson. 2016. Lipolytic and thermogenic depletion of adipose tissue in cancer cachexia. Semin Cell Dev Biol 54:68–81.

Tsujinaka, T., J. Fujita, C. Ebisui, M. Yano, E. Kominami, K. Suzuki, K. Tanaka, A. Katsume, Y. Ohsugi, H. Shiozaki, and M. Monden. 1996. Interleukin 6 receptor antibody inhibits muscle atrophy and modulates proteolytic systems in interleukin 6 transgenic mice. J Clin Invest 97:244–249.

van Hall, G. 2012. Cytokines: muscle protein and amino acid metabolism. Curr Opin Clin Nutr Metab Care 15:85–91.

von Haehling, S., M.S. Anker, and S.D. Anker. 2016. Prevalence and clinical impact of cachexia in chronic illness in Europe, USA, and Japan: facts and numbers update 2016. J Cachexia Sarcopenia Muscle 7:507–509.

Washington, T.A., J.P. White, J.M. Davis, L.B. Wilson, L.L. Lowe, S. Sato, and J.A. Carson. 2011. Skeletal muscle mass recovery from atrophy in IL-6 knockout mice. Acta Physiol (Oxf) 202:657–669.

Zhang, Y., W. Yan, M.A. Collins, F. Bednar, S. Rakshit, B.R. Zetter, B.Z. Stanger, I. Chung, A.D. Rhim, and M.P. di Magliano. 2013. Interleukin-6 is required for pancreatic cancer progression by promoting MAPK signaling activation and oxidative stress resistance. Cancer Res 73:6359–6374.

